# PAG1 directs SRC-family kinase intracellular localization to mediate receptor tyrosine kinase-induced differentiation

**DOI:** 10.1101/2020.02.20.958496

**Authors:** Lauren Foltz, Juan Palacios-Moreno, Makenzie Mayfield, Shelby Kinch, Jordan Dillon, Jed Syrenne, Tyler Levy, Mark Grimes

**Affiliations:** Division of Biological Sciences, Center for Biomolecular Structure and Dynamics, and Center for Structural and Functional Neuroscience, The University of Montana, Missoula MT 59812, USA; University of Minnesota Medical School, Duluth MN 55812, USA; Cell Signaling Technology, Danvers, MA 01923, USA

## Abstract

All receptor tyrosine kinases (RTKs) activate similar downstream signaling pathways through a common set of effectors, yet it is not fully understood how different receptors elicit distinct cellular responses to cause cell proliferation, differentiation, or other cell fates. We tested the hypothesis that regulation of SRC Family Kinase (SFK) signaling by the scaffold protein, PAG1, influences cell fate decisions following RTK activation. We generated a neuroblastoma cell line expressing a PAG1 fragment that lacks the membrane spanning domain (PAG1™^-^) and localized to the cytoplasm. PAG1™^-^ cells exhibited higher amounts of active SFKs and increased growth rate. PAG1™^-^ cells were unresponsive to TRKA and RET signaling, two RTKs that induce neuronal differentiation, but retained responses to EGFR and KIT. Under differentiation conditions, PAG1™^-^ cells continued to proliferate and did not extend neurites or increase β-III tubulin expression. FYN and LYN were sequestered in multivesicular bodies (MVBs), and dramatically more FYN and LYN were in the lumen of MVBs in PAG1™^-^ cells. In particular, activated FYN was sequestered in PAG1™^-^ cells, suggesting that disruption of FYN localization led to the observed defects in differentiation. The results demonstrate that PAG1 directs SFK intracellular localization to control activity and to mediate signaling by RTKs that induce neuronal differentiation.

## Introduction

Precise temporal and spatial control over cell signaling pathways is necessary to coordinate diverse cell responses to extracellular signals (Bergeron et al., 2016; Irannejad et al., 2015; Lemmon et al., 2016). RTKs initiate intracellular signaling pathways after binding extracellular ligands at the plasma membrane, which typically induces receptor homodimerization and transphosphorylation of tyrosine residues on the cytoplasmic tail of dimer partners (Lemmon and Schlessinger, 2010). Phosphorylated tyrosine residues on RTKs act as platforms for the recruitment of adaptor proteins to activate four downstream canonical cell signaling pathways: RAS/MAPK, PI3K, PLC-γ, and SFKs. Different RTKs elicit different cell responses, however, and not all RTKs activate each pathway to the same extent. How different receptors initiate and terminate a cascade of effector signals with precise sequence and timing to elicit a particular cell response remains unclear.

The SFK non-receptor tyrosine kinases comprise nine members in humans: SRC, YES1, FYN, FGR, FRK, LYN, BLK, HCK, and LCK. SFKs have a modular domain structure and are composed of a unique N-terminal region termed the SRC Homology 4 domain (SH4), an SH3 domain, an SH2 domain, an SH1 catalytic/kinase domain and a short regulatory tail. SH2 and SH3 domains bind to phosphorylated tyrosine residues and proline rich regions, respectively (Mayer, 2015). This domain structure, similar to SH2/SH3 adaptor proteins, links SFKs to a large interactome tuned to phosphotyrosine signals, propagated through their tyrosine kinase activity, which is governed by both activating and inhibitory phosphorylation (Harrison, 2003; Pawson, 2004; Sandilands and Frame, 2008). Myristoylation and palmitoylation on the N-terminal region recruits SFKs to membrane surfaces, including the plasma membrane, endosomes, Golgi, and endoplasmic reticulum, although a pool of SFKs remains cytosolic (Alland et al., 1994; Hantschel et al., 2003). SFKs are thus signaling hubs whose activity can be placed on a variety of intracellular membranes to link upstream RTK signaling to the appropriate effector pathways.

Our previous work pointed to the dynamic intracellular localization of two members of the SRC family of tyrosine kinases, FYN and LYN, as potential means to control RTK effector pathways in cells from the neural crest-derived cancer, neuroblastoma. In a phosphoproteomic study, we combined fractionation of membrane compartments including endosomes and detergent-resistant lipid rafts (McCaffrey et al., 2009; Pryor et al., 2012) with large scale analysis of 21 neuroblastoma cell lines (Palacios-Moreno et al., 2015). Neuroblastoma cells express more than half of the RTKs in the human genome, which suggests that mechanisms to discern different receptors’ signals, and integrate them when more than one receptor is activated, must play a role in cell fate decisions in neural crest and neuroblastoma, as for other tissues in multicellular organisms (Chen et al., 2012a; Iglesias-Bartolome and Gutkind, 2011; Schwarz et al., 2012; Theveneau and Mayor, 2012). Different RTKs were enriched in endosomes and lipid rafts in different cell lines (Palacios-Moreno et al., 2015). Similarly, SFK family members, mainly FYN and LYN (and to a lesser extent SRC and YES1), phosphorylated on both activating and inhibitory sites, were enriched in different endosome populations and plasma membrane fractions with distinct distributions in different neuroblastoma cell lines (Palacios-Moreno et al., 2015). FYN and LYN activity and localization change in response to activation from different RTKs, which suggests that a mechanism must exist to tailor SFK relocalization to elicit a particular response (Palacios-Moreno et al., 2015).

We also observed enrichment of PAG1 (Phosphoprotein Associated with Glycosphingolipid enriched microdomains 1) in both endosomes and lipid rafts (Palacios-Moreno et al., 2015). PAG1 is a scaffold protein composed of a very short extracellular N-terminal domain, a transmembrane domain, and a long intracellular cytoplasmic domain. PAG1’s cytoplasmic tail is rich in phosphorylation sites and potential docking sites for interactions with SFKs, phosphatases, and trafficking machinery (Barua et al., 2012). PAG1 binds SFKs and regulates their activity by facilitating the interaction with C-terminal SRC Kinase (CSK) (Ingley et al., 2006; Kawabuchi et al., 2000; Lindquist et al., 2011; Oneyama et al., 2008). Docking of both CSK and SFKs on PAG1 brings them in proximity where CSK phosphorylates SFKs on C-terminal inhibitory tyrosine residues (Lindquist et al., 2003; Saitou et al., 2014; Solheim et al., 2008a). Due to its known functions and presence in lipid rafts and endosomes, we hypothesized that PAG1 may direct SFK intracellular location as well as activity, and that this mechanism may play a role in distinguishing cell signaling responses to different receptors.

Here we investigate cell signaling mechanisms that distinguish RTK signals that cause proliferation vs. differentiation in neuroblastoma cell lines. We show that generation of a neuroblastoma cell line expressing a PAG1 mutant lacking the transmembrane domain (PAG1™^-^) resulted in enhanced tumorigenic phenotypes: anchorage-independent growth and proliferation. PAG1™^-^-expressing cells failed to respond to differentiative signaling, while maintaining responses to oncogenic receptors. We show that cells sequestered FYN and LYN in the lumen of multivesicular bodies (MVBs), and that this process was disrupted in PAG1™^-^ cells, which exhibited high amounts of active FYN, and inactive LYN, sequestered in MVBs. Our findings show that PAG1 regulates the intracellular localization and activity of FYN and LYN to convey signaling instructions from RTKs that cause neuronal differentiation.

## Results

To examine the role of PAG1 in SFK regulation, we aimed to generate human neuroblastoma PAG1 knockout cell lines using lentiviral delivery of CRISPR/Cas9 plasmids. Sequencing results after transduction and selection of SH-SY5Y cells indicated a heterozygous frame shift mutation in Exon 5 of the PAG1 gene that resulted in a stop codon shortly downstream (Figure 1A, B). We expected reduced expression, however, a mutant PAG1 protein that lacks the transmembrane domain was detected by western blot and mass spectrometry analysis (PAG1™^-^, Figure 1C). This protein was not bound to membranes, was absent in endosomes, and strictly located to cytosolic fractions (Figure 1D). WT PAG1 was localized in endosome fractions that migrated to the lower half of the gradient (Figure 1D). The data suggest that the soluble, cytoplasmic PAG1™^-^ protein was translated starting from one of two downstream methionine residues, which have Kozak consensus sequences and predicted protein products that are truncations of the wild-type protein (Mou et al., 2017; Sharpe and Cooper, 2017; Wang et al., 2002). All PAG1 peptides identified by mass spectrometry lie within the range of the predicted truncated protein.

**Figure 1:**
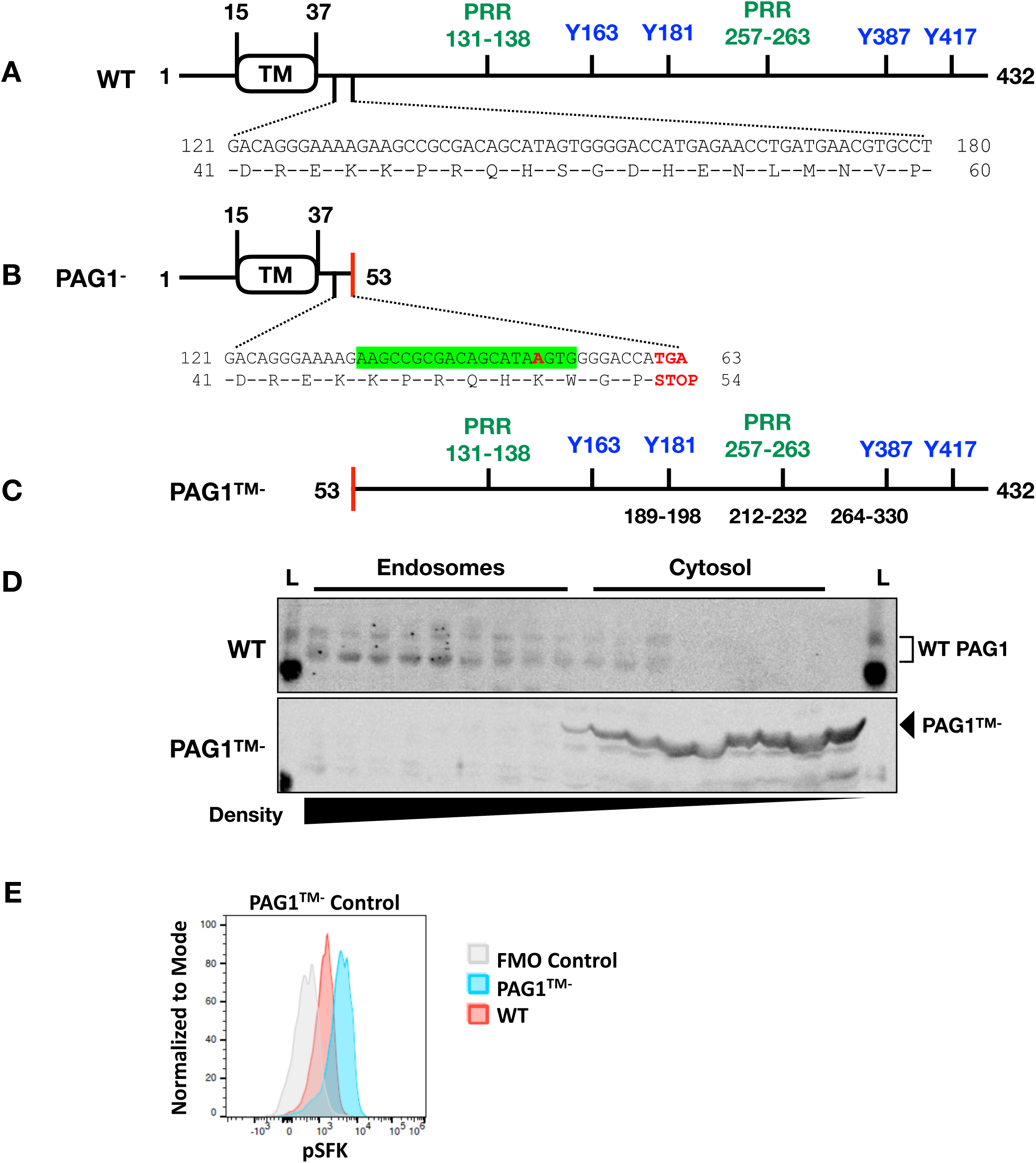
SH-SY5Y cells expressing cytosolic PAG1 (PAG1™^-^) were generated using CRISPR/Cas9. (A) Wild type (WT) PAG1 has a small extracellular N-terminal region, transmembrane domain (TM), and long intracellular C-terminal tail. Several phosphorylated residues and proline rich regions (PRR, PXXP motif, green) act as docking sites for protein-protein interactions. Phosphorylated Y163/Y181 and Y387/417 (blue) are binding sites for FYN and LYN SH2 domains, respectively (Barua et al., 2012; Ingley et al., 2006; Solheim et al., 2008a, 2008b). (B) CRISPR sgRNA recognition site is highlighted in green. Sequencing results indicated a single base pair insertion that resulted in a stop codon to generate a heterozygous PAG1^-^ SH-SY5Y cell line (red). (C) Met residues upstream of the detected peptides act as alternate translation start sites to produce the cytosolic PAG1 product (PAG1™^-^). PAG1 peptides were detected via mass spectrometry on PAG1^-^ cell lines (amino acid numbers indicated in black). (D) Organelle fractionation comparing the endosomal distribution of WT PAG1 vs. PAG1™^-^. Fractions were taken from 25% to 2.5% iodixanol gradients and decrease in density from left to right. L= biotinylated ladder. (E) Levels of activated SFKs were increased in PAG1™^-^ expressing SH-SY5Ys (blue) compared to wild type SH-SY5Y cells (red). Cells were stained with a pSFK antibody recognizing the activating phosphorylation site conserved in several SFKs and pSFK fluorescence was measured by flow cytometry.

We hypothesized that PAG1™^-^-expressing cells would have disrupted regulation of SFKs and other signaling proteins downstream of RTK signaling. To test this, we determined the SFK activation status of individual neuroblastoma cells using flow cytometry. We assessed total SFK phosphorylation using an antibody that recognizes the activating phosphorylation site (pY416), and found that the PAG1™^-^ cell population had nearly twice the amount of active SFKs compared to wild type cells (Figure 1E), suggesting that the mutant PAG1 is deficient in its ability to control SFK activity. The generation of PAG1™^-^-expressing cells presented a unique opportunity to study the effect of dysregulated SFK activity on cell growth, differentiation, and cell signaling pathways, without directly affecting SFK structure or binding capabilities.

### PAG1™^-^ enhanced growth rate in neuroblastoma cells and reduced the effect of the SFK inhibitor PP2

To compare the growth of PAG1™^-^ and wild type SH-SY5Y cells, we employed the MTT assay, which determines cell growth by measuring mitochondrial enzyme activity. Because growth rate measurements have been shown to fluctuate, even within the same cell line, we used a more robust method described by Hafner et al. to calculate growth rate inhibition induced by drug treatment (Hafner et al., 2016). To further characterize changes in proliferation rate due to overactive SFK signaling, we treated cells with a small molecule SFK inhibitor, PP2. PP2 is very active against LCK and FYN, but less active against SRC and LYN (Brandvold et al., 2012; Hanke et al., 1996). Figure 2A compares the growth rate of WT cells treated with PP2 to PAG1™^-^ cells treated with PP2 to determine the effect of the PAG1™^-^ truncation. In the equation described by Hafner et al., GRI = 2^(R’/R)^ – 1, where R = WT growth rate and R’ = PAG1™^-^ growth rate (see complete formulae in methods). A GRI of 3.17 clearly indicates that PAG1™^-^ cells grow faster than WT cells (Figure 2A, grey). The increase in GRI with increasing PP2 concentration (Figure 2A, green) suggests that either WT cells are more sensitive to PP2 compared to PAG1™^-^ cells, or that PAG1™^-^ cells increase their growth rate (Figure 2A). To distinguish these possibilities, the growth rate of WT and PAG1™^-^ cells were compared to their respective vehicle controls (Figure 2B). The data indicate that PP2 did not greatly affect the growth rate of PAG1™^-^ cells, but did significantly reduce WT cell growth. Since PAG1™^-^ cells had increased SFK activity and were less sensitive to the SFK inhibitor PP2 than WT cells, this suggests that disruption of PAG1 function probably affects more than one SFK, and that PP2-sensitive SFKs are not driving cell proliferation in PAG1™^-^ cells.

**Figure 2:**
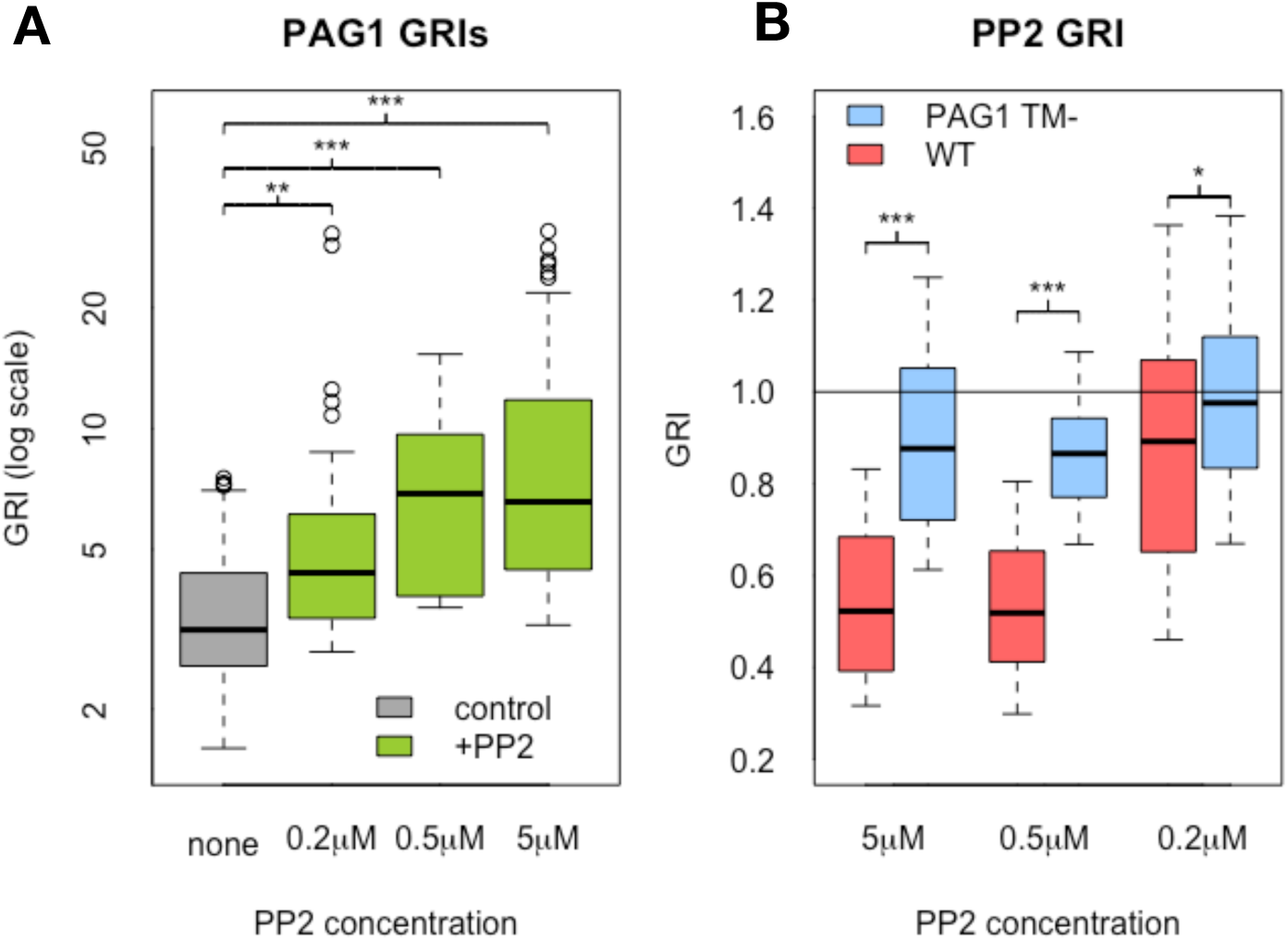
PAG1™^-^ expression enhanced growth rate and de-sensitized cells to the SFK inhibitor PP2. (A and B) MTT analysis of WT and PAG1™^-^ growth rates with PP2 (0.2M, 0.5M, and 5M) addition. Growth rate index (GRI) was calculated as described in Hafner et al. (2016; described in Methods). In A, the growth rate ratio compares the growth of wild type cells to PAG1™^-^ cells under control (grey) and drug-treated conditions (green). In B, the growth rate ratio compares the growth of WT and PAG1™^-^ cells with and without PP2. *P<0.05, **P<0.005, ***P<0.0005. Student’s t test was also calculated for pooled growth rates of WT vs. PAG1™^-^ cells.

### PAG1™^-^ cells exhibited increased anchorage-independent growth

We next asked whether PAG1™^-^ expression contributed to the gain of transformed tumorigenic phenotypes as measured by colony growth in soft agar. PAG1™^-^ cells exhibited increased colony formation in soft agar compared to wild type cells (Figure 3A, B). Cells expressing PAG1™^-^ formed more total colonies than wild type cells, and PAG1™^-^ colonies were much larger, consistent with the increased cell division noted above. PP2 treatment did not significantly affect colony formation for PAG1™^-^ expressing cells, but did decrease the number and size of colonies formed by WT cells (Figure 3A). These findings are consistent with previously reported experiments using siRNA knockdown of PAG1 (Agarwal et al., 2016; Oneyama et al., 2008).

**Figure 3:**
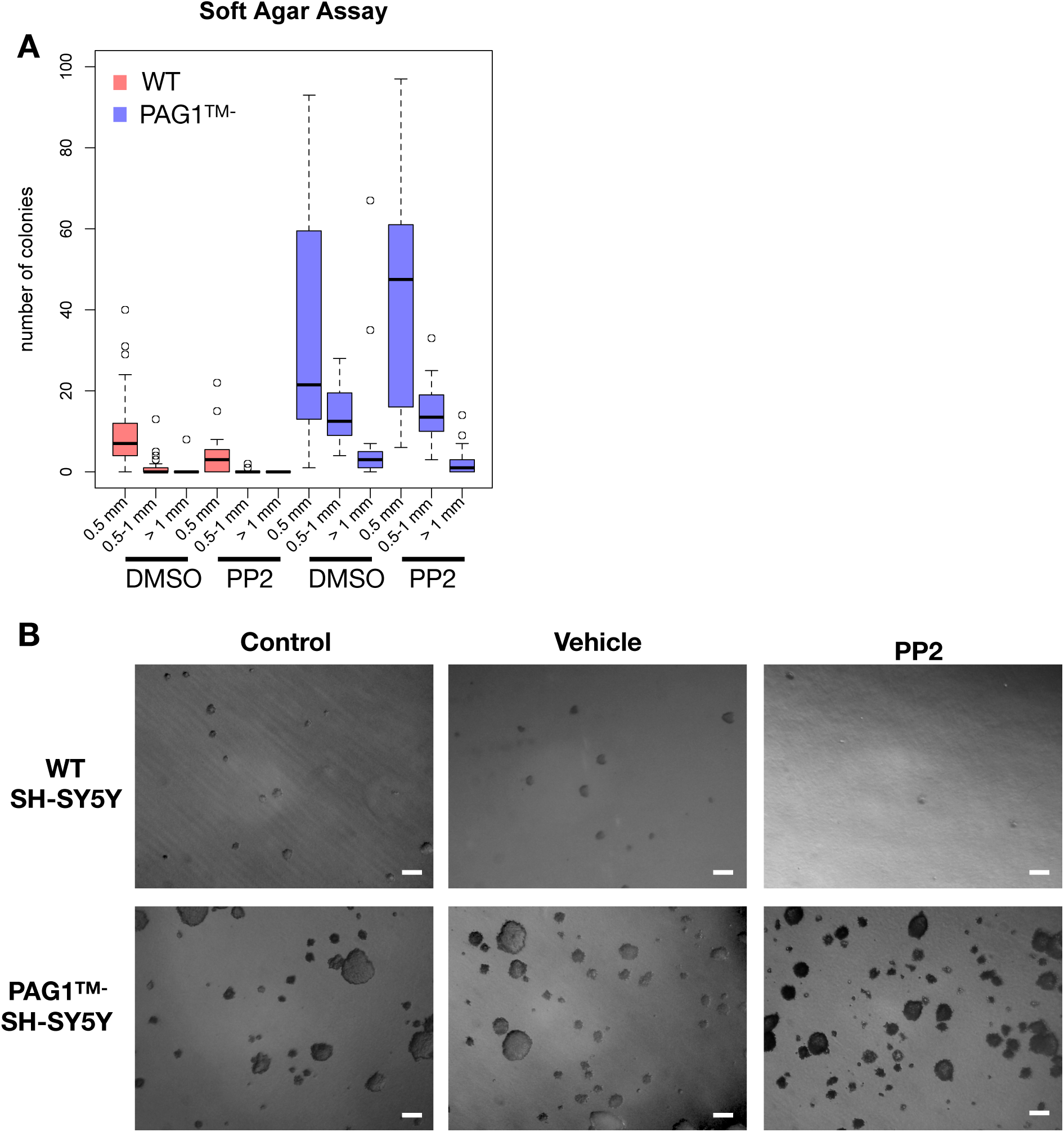
PAG1™^-^ enhanced anchorage-independent growth. (A) Soft agar assay of wild type vs. PAG1™^-^ expressing SH-SY5Y cells. After 7 days, control or PP2 treated (5µM) cells were imaged and colony sizes were binned into three categories (0.5mm^2^, 0.5-1mm^2^, and 1mm^2^). Data are a summary of three independent experiments. (B) Representative images of colonies quantified in A for each condition. Scale bar = 1mm.

### PAG1™^-^ prevented differentiation of SH-SY5Y neuroblastoma cells

Different RTKs induce distinct cell fate decisions that are mediated by SFK signaling and other pathways. Because increases in tumorigenicity and proliferation are typically accompanied by deficits in differentiation, we hypothesized that disrupting SFK activation by expression of the PAG1™^-^ mutant would also disrupt differentiation. We used neurite extension and expression of β-III tubulin as assays for differentiation. We measured neurite length after exposing cells to a combination of retinoic acid (RA) and NGF, which induces neuronal differentiation in neuroblastoma cell lines (Shipley et al., 2016). SH-SY5Y cells normally respond to these ligands by slowing cell division and extending neurites (Figure 4A, B). PAG1™^-^-expressing cells did not extend neurites under differentiation conditions, whereas control SH-SY5Ys displayed significant increases in neurite length (Figure 4A). Similarly, we assessed cellular levels of β-III tubulin by flow cytometry after growth in differentiation inducing media. β-III tubulin is a neuron-specific tubulin whose expression is induced by neuronal differentiation (Figure 4C; Guo et al., 2010). PAG1™^-^ cells did not increase β-III tubulin expression after exposure to differentiation conditions (blue), whereas wild-type cells displayed robust induction of expression (red; Figure 4C). The normal slowing of the cell division cycle upon differentiation was also disrupted in mutant cells. While the majority of wild type cells remained in G0/G1, PAG1™^-^ cells had higher percentages of cells in S and G2/M stages of the cell cycle while exposed to these differentiation conditions, suggesting frequent cell division (Figure 4D, E). These results suggest that PAG1™^-^ SH-SY5Ys are not capable of inducing cell signaling mechanisms that promote neuronal differentiation.

**Figure 4:**
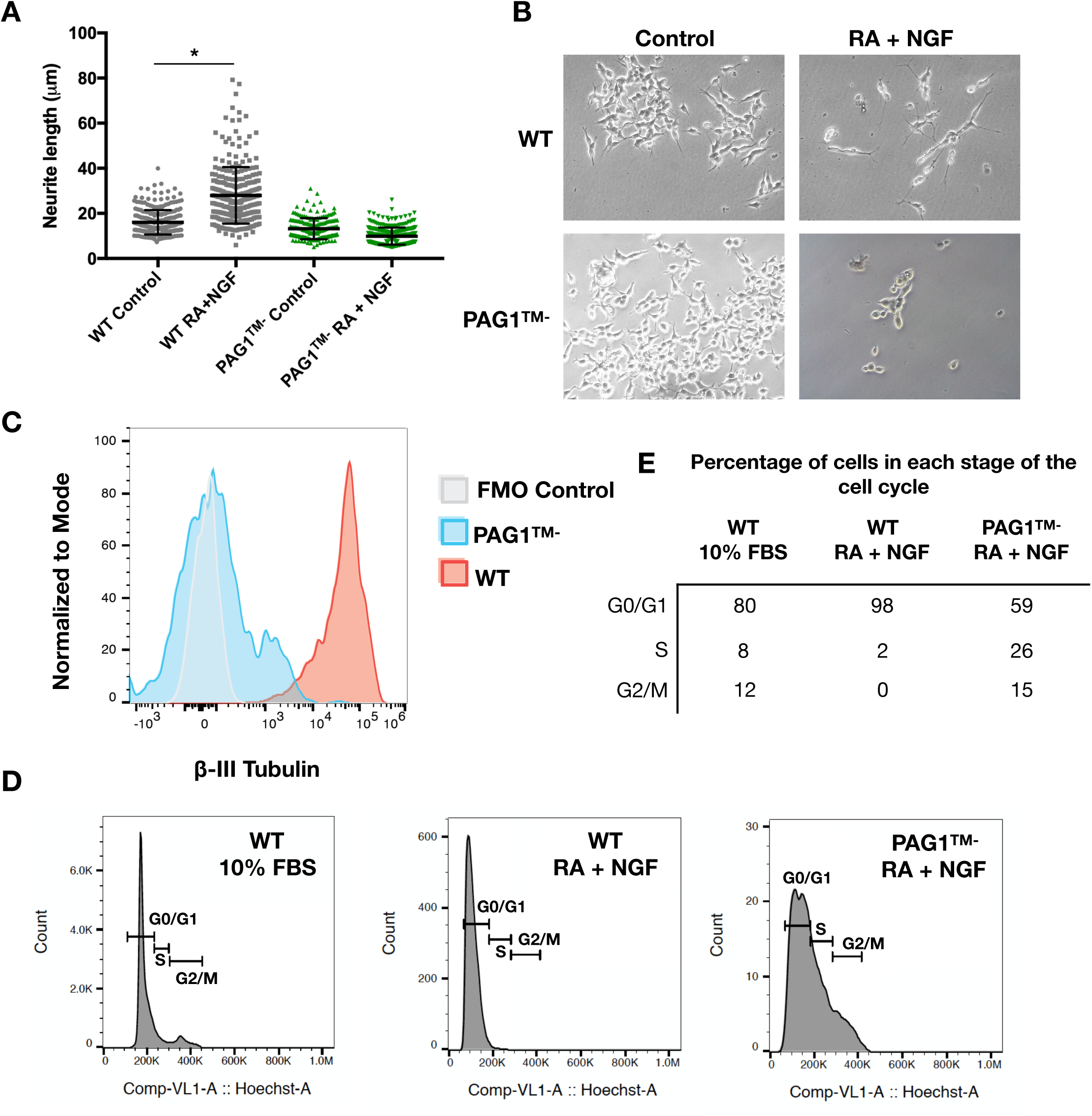
PAG1™^-^ prevented differentiation in SH-SY5Y cells. (A) Neurite lengths of wild type and PAG1™^-^ SH-SY5Y cells after growth in control conditions (RPMI 1640, 2% FBS) and in differentiation conditions (RPMI 1640, 2% FBS, 10µM Retinoic Acid (RA), 5nM nerve growth factor (NGF)) * = p<0.05 (B) Representative images of neurites after 8 days of growth in the indicated conditions, 20X magnification. (C) Flow cytometry of β-III tubulin expression, a marker of neuronal differentiation. (D) Cell cycle analysis of wild type SH-SY5Y and SH-SY5Y PAG1™^-^ cells by flow cytometry. Cells were seeded in standard growth medium (RPMI 1640, 10% FBS) on collagen coated plates, and were exposed for 96 hrs. to 10µm RA and 5nM NGF in low serum media (2% FBS). Cells were then stained with Hoechst 33342 and relative DNA content was measured by flow cytometry. (E) The percentage of cells in each stage of the cell cycle for each condition in D.

### PAG1™^-^ expression increased ERK activation in response to EGF

Because PAG1™^-^ cells exhibited enhanced growth rate and defects in differentiation, we hypothesized that downstream cell signaling responses to different RTKs would reflect these characteristics. We asked whether changes in SFK signaling by PAG1™^-^ expression affected the activation of the RAS/MAPK pathway. We assessed the activation of ERK and SFKs for both wild type and PAG1™^-^ expressing SH-SY5Y neuroblastoma cells after 5 and 60-minute stimulations with different RTK ligands.

While activation of EGFR induced a robust pERK response in both cell types, PAG1™^-^ cells had significantly more activated ERK, especially after 5 and 60 minutes of EGF stimulation (Figure 5A, B). Treatment with EGF caused a modest increase in pSFK activation in wild-type cells after both 5 and 60 minutes (Figure 5C, D). PAG1™^-^ cells started at a higher baseline of pSFK activation (Figure 1E), and there was no further increase in the amount of active SFKs after exposure to EGF. These results suggest that PAG1™^-^ cells retained their ability to activate the RAS/MAPK cell signaling pathway in response to EGF despite a high baseline of activated SFKs.

**Figure 5:**
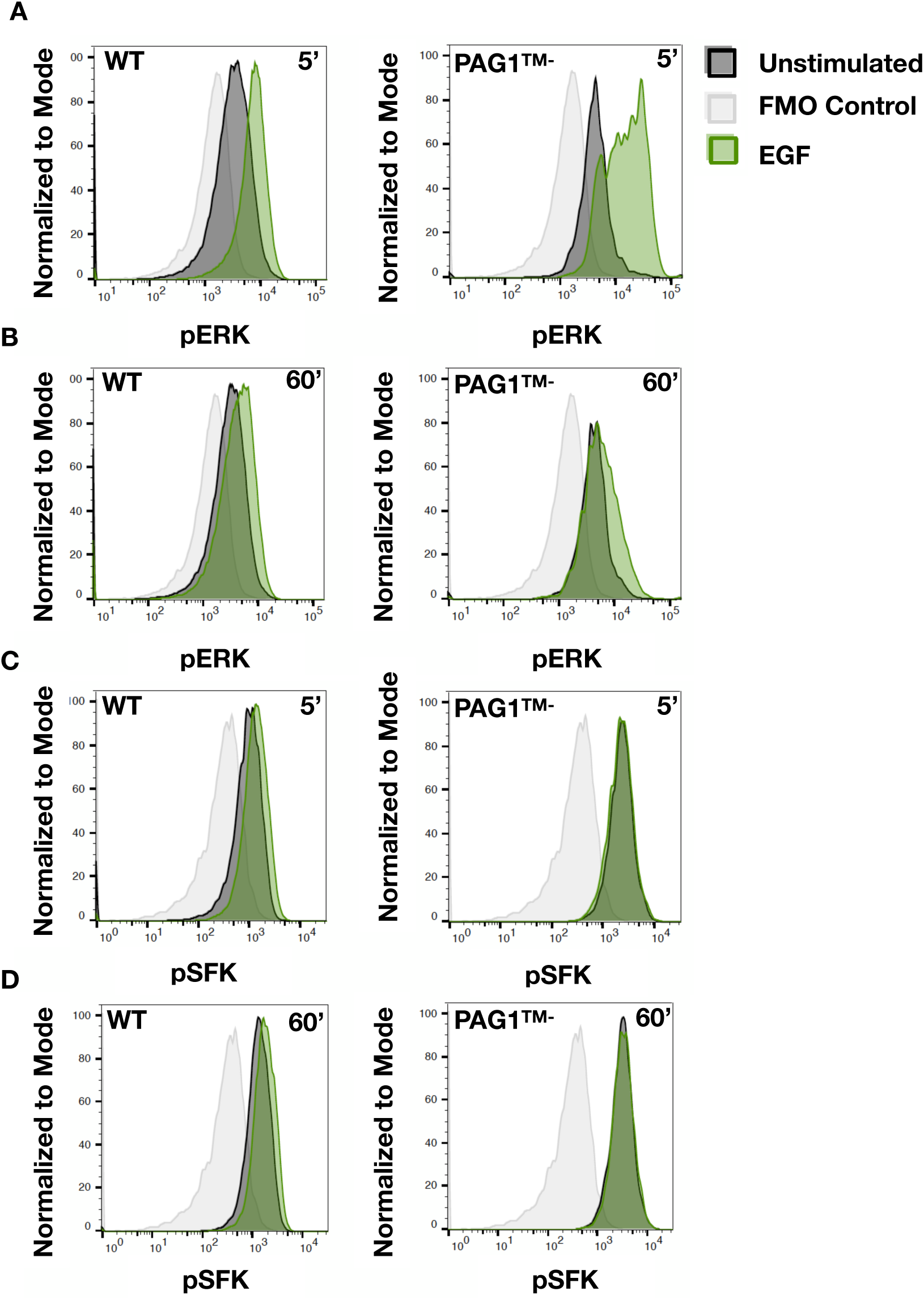
PAG1™^-^ enhanced the pERK response to EGF. Flow cytometry analysis of pERK (A, B) and pSFK (C, D) activation after exposure to 5nM EGF for either 5 minutes (A and C) or 60 minutes (B and D). Adherent SH-SY5Y cells expressing either wild-type (WT) PAG1 or PAG1™^-^ were treated with ligand before fixing and staining with fluorescent antibodies for pERK (Phospho-p44/42 MAPK (Erk1/2) (Thr202/Tyr204)) and pSFKs (pY416). Data displayed are from one experiment representative of at least three independent experiments. Cell counts were normalized to mode to account for differences in final cell number.

### PAG1™^-^ increased SRC activation in response to SCF

We then asked if the signaling responses observed after EGFR stimulation were similar to those of another typically pro-oncogenic receptor, KIT. We had previously shown that KIT stimulation by SCF caused an increase of SFKs in endosomes in LAN-6 neuroblastoma cells (Palacios-Moreno et al., 2015), so we asked whether expression of PAG1™^-^ would affect ERK and SFK responses. KIT activation did not induce a pERK response in either wild type or PAG1™^-^ cells (Figure 6A, B). However, SCF treatment did increase total pSFK levels at 5 minutes in both cell types (Figure 6C, D). As noted above, PAG1™^-^ cells started at higher basal levels of pSFK activation, but with SCF the amount of pSFKs increased further upon ligand stimulation, mainly at 5 minutes. Wild type cells started at a low level of pSFK activation but exhibited a similar increase in pSFK levels upon ligand stimulation. These results show that PAG1™^-^ cells retain the ability to respond to SCF by further increasing SFK activation.

**Figure 6:**
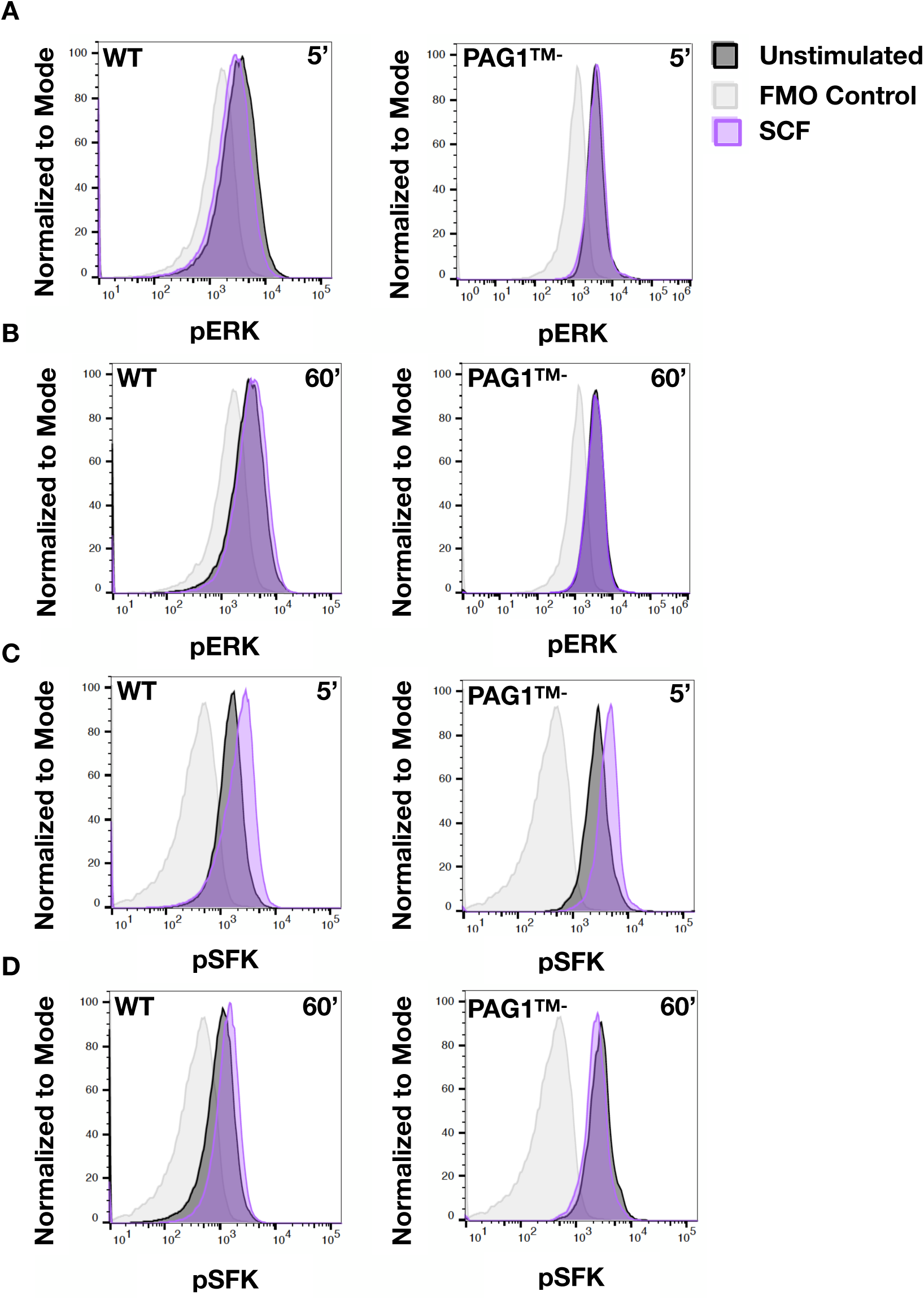
PAG1™^-^ increased SRC response to KIT activation. Flow cytometry analysis of pERK and pSFK activation as in Figure 5 except cells were exposed to 5nM SCF for either 5 minutes or 60 minutes. Data displayed are from one experiment representative of at least three independent experiments.

### PAG1™^-^ abrogated ERK activation in response to NGF and GDNF

That PAG1™^-^ cells failed to differentiate (Figure 4) suggests that cell signaling mechanisms activated in response to RTKs that induce neuronal differentiation may be impaired. Stimulation of TRKA (NTRK1) and RET by NGF and GDNF, respectively, is known to induce differentiation and survival in neurons and neuroblastoma cells (Harrington et al., 2011; Miller and Kaplan, 2001). While wild type cells exhibited a robust increase in pERK in response to NGF, the pERK response was entirely absent in PAG1™^-^ cells (Figure 7A, B). TRKA activation did lead to small increases in pSFK activation in both wild type and PAG1™^-^ cells, however this response was short-lived and largely gone by sixty minutes (Figure 7C, D). Activation of RET by GDNF induced similar pERK responses in wild type, but not PAG1™^-^ cells, and GDNF induced a more robust early pSFK stimulation in wild type but not PAG1™^-^ cells (Supplementary Figure 1). These results show that PAG1™^-^ cells are unable to activate the RAS/MAPK pathway in response to two different RTKs that promote neuronal differentiation.

**Figure 7:**
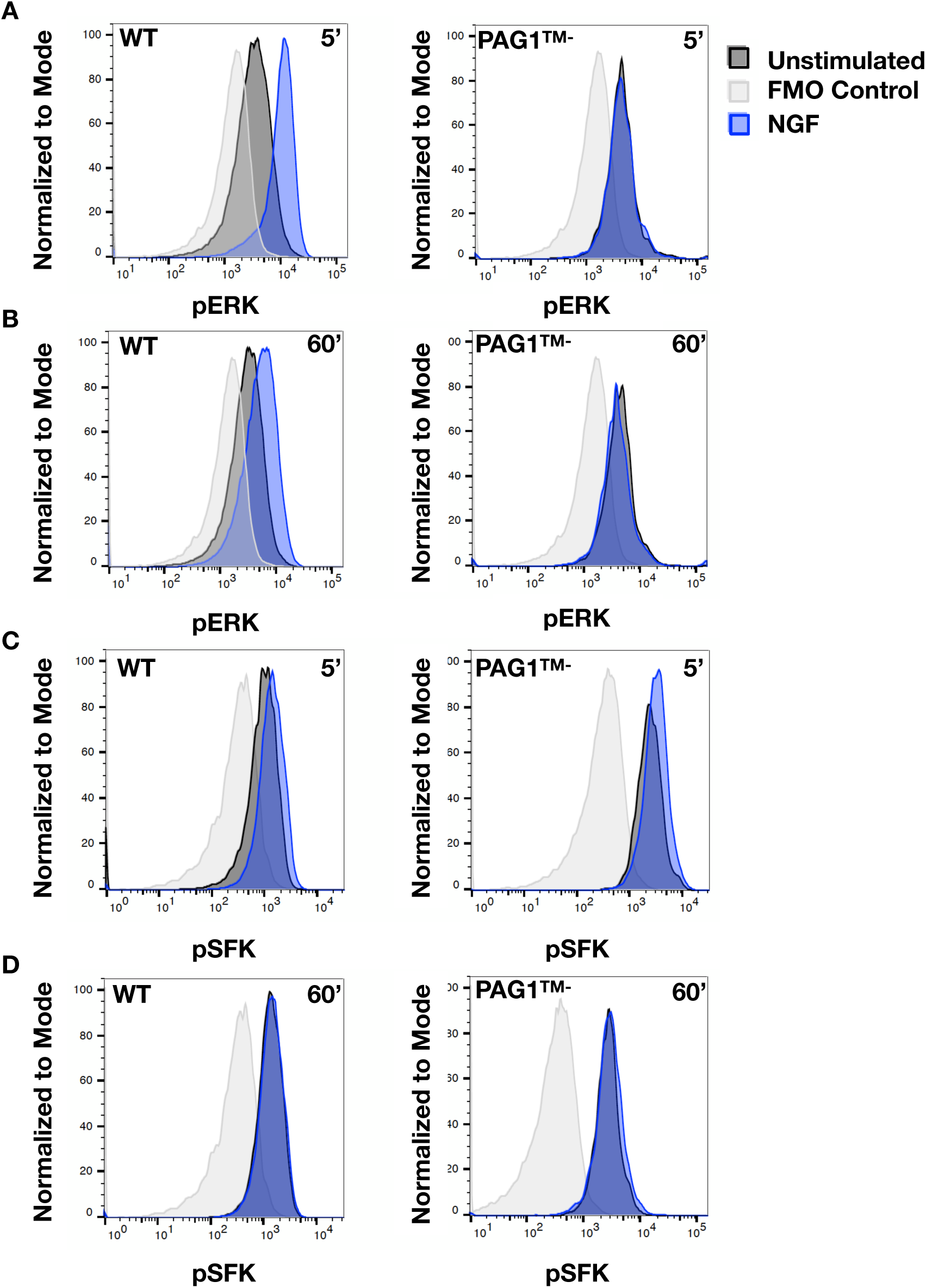
PAG1™^-^ decreased responses to pro-differentiation receptor TRKA. Flow cytometry analysis of pERK and pSFK activation as in Figure 5 except cells were exposed to 5nM NGF for either 5 minutes or 60 minutes. Data displayed are from one experiment representative of at least three independent experiments.

The AKT/PI3K pathway is also activated by RTK signaling; however, we did not detect any changes in AKT phosphorylation that were not associated with the G2/M stage of the cell cycle (Supplementary Figure 2A). PAG1™^-^ cells had a higher growth rate (Figure 2) and greater percentage of cells in G2/M, which likely explains a greater percentage of cells with high pAKT (pT308) compared to WT cells (Supplementary Figure 2B).

These results distinguish downstream cell signaling responses to different RTKs and indicate that PAG1 is necessary for cells to respond to RTKs that promote neuronal differentiation. Responses to proliferative signals remain intact in the presence of cytoplasmic truncated PAG1™^-^, which we hypothesize acts as a dominant negative by binding regulatory proteins such as CSK in the cytosol and keeping them from interacting with SFKs on membranes. In the canonical view, KIT, EGFR, NGF, and RET should all activate the RAS/MAPK and SFK pathways in response to ligand, yet these receptors produce different responses. Our results so far suggest that there is an additional layer of regulation between receptor activation and ERK/SFK activation. We hypothesized that PAG1’s role is to control the activity and intracellular localization of members of the SFK family. We focus on FYN and LYN because of their robust expression in neuroblastoma cell lines and dynamic localization to endosomes and other cell membranes (Palacios-Moreno et al., 2015).

### PAG1™^-^ expression altered FYN and LYN distribution in endosomes

Because PAG1 was detected in endosomes with FYN and LYN (Palacios-Moreno et al., 2015 b61), we hypothesized that PAG1™^-^ expression may affect the endosomal localization of FYN and LYN. To determine SFK localization in wild type and PAG1™^-^ expressing cells, we performed organelle fractionation experiments to isolate endosomes as previously described (McCaffrey et al., 2009; Palacios-Moreno et al., 2015). Under conditions of serum starvation, the distribution of FYN and LYN in endocytic organelles was distinct in both PAG1™^-^ and wild-type cells (Supplementary Figure 3), consistent with data from LAN-6 neuroblastoma cells (Palacios-Moreno et al., 2015). When KIT was activated by SCF, more FYN and LYN were detected in endosomal fractions of PAG1™^-^ cells. These effects were similar to those observed previously, though ligand-induced redistribution of FYN and LYN was not as dramatic as in LAN-6 cells (Palacios-Moreno et al., 2015).

### FYN and LYN were sequestered in the lumen of multivesicular bodies (MVBs), and their sequestration was affected by PAG1™^-^ expression

Taelman et al. first reported sequestration of the kinase GSK3β in intraluminal vesicles of multivesicular bodies (MVBs) as a means of controlling kinase access to cytoplasmic substrates (Taelman et al., 2010; Vinyoles et al., 2014). We hypothesized that PAG1™^-^ may regulate FYN and LYN access to cytosolic substrates by sequestration into MVBs in a similar manner. To test this, we isolated endosome fractions via organelle fractionation and performed a protease protection assay. To examine the effects of proliferative signals, cells were either grown in 10% FBS containing medium (+serum) or serum starved (-serum) for two hours.

After thirty minutes treatment with Protease K, both FYN and LYN were protected in endosome fractions, but degraded in the presence of detergent, indicating that these proteins reside in the lumen of membrane-enclosed organelles such as MVBs (Figure 8A). The lower molecular weight isoform of FYN, FYN isoform C, was preferentially protected (Figure 8A, <60kD band). Transferrin receptor, a marker of recycling endosomes, exhibited minimal protection after five minutes of incubation with protease, and was not protected after thirty minutes of protease treatment, as expected for a protein that is not in the lumen of MVBs. We also detected FYN and LYN (but not TfR) in exosomes isolated from WT and PAG1™^-^ cell culture media (Figure 8B). These data strongly suggest that localization of both FYN and LYN in the lumen of MVBs is a means to control their activity by limiting access to cytosolic substrates.

**Figure 8:**
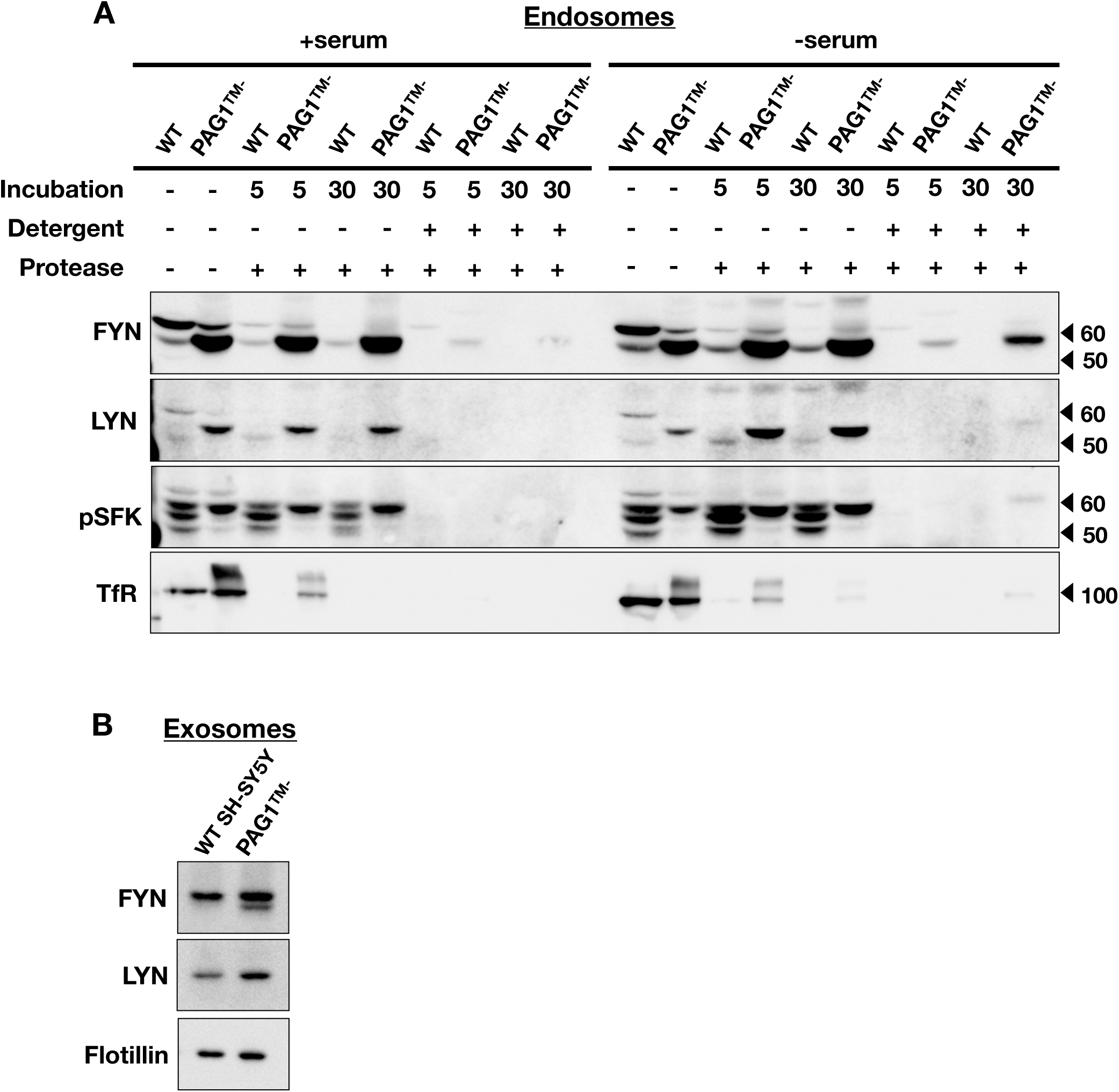
Wild-type and PAG1™^-^ SH-SY5Ys sequestered active SFKs in multivesicular bodies. (A) SH-SY5Y cells expressing either WT PAG1 or PAG1™^-^ were grown in either 10% FBS containing medium (+serum) or serum starved (-serum) before cellular fractionation. The endosome fraction was equally distributed into five parts before treatment with the indicated combinations of detergent [0.1% IGEPAL] and/or Protease K [0.01µg/µL]. Treatments were incubated for 5 minutes (5) or 30 minutes (30) at 37°C and stopped by TCA addition. Transferrin receptor (TfR), a marker for recycling endosomes, was used as a negative control for sequestration. (B) Exosomes isolated from WT SH-SY5Y or PAG1™^-^ SH-SY5Y growth mediums. Western blots show FYN, LYN, and flotillin in exosomes; we detected no transferrin receptor in exosomes (not shown).

There were several differences between WT and PAG1™^-^ cells. First, there was more total FYN and (to a lesser extent) LYN sequestered in the lumen of MVBs from PAG1™^-^ cells, indicated by protease protection and presence in exosomes (Figure 8). Similar amounts of endosomes were recovered from each cell type in these experiments indicated by TfR (Figure 8A, 9A, TfR). Second, while the amounts of activated SFKs protected from protease were somewhat similar, WT cells had several pSFKs, whereas PAG1™^-^ cells predominantly had a single activated SFK (pSFK, Figure 8A). Comparing pSFKs to total FYN and LYN, the data indicate that WT cells preferentially sequestered activated SFKs, and PAG1™^-^ cells sequestered more total FYN and LYN (so the ratio of active to total SFKs was lower). FYN was predominantly recovered by immunoprecipitation with anti-pSFK antibodies in endosome fractions in PAG1™^-^ cells (Figure 9A), indicating that the single pSFK band in PAG1™^-^ cells in Figure 8A is FYN. Much less LYN was recovered by immunoprecipitation from PAG1™^-^ endosomes (Figure 9A).

**Figure 9:**
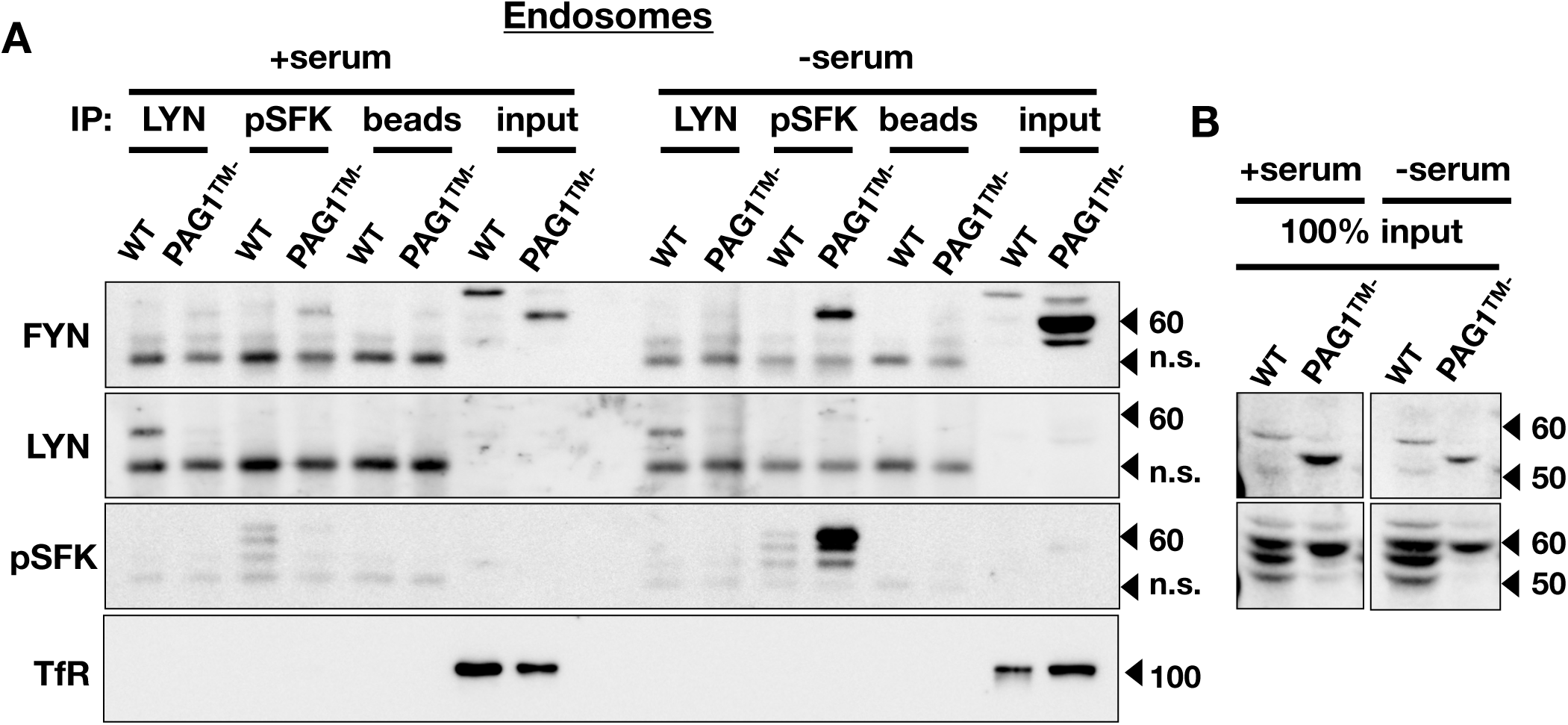
FYN was the major active SFK in PAG1™^-^ SH-SY5Y cells. LYN and pSFKs (Y416) were immunoprecipitated (IP) from endosome fractions of WT and PAG1™^-^ SH-SY5Y cells grown in either 10% FBS containing medium (+serum) or serum starved (-serum). Fractions were equally distributed into four parts for immunoprecipitations. Samples were run on SDS-PAGE and the membrane was probed for LYN, FYN, pSFK (Y416), and Transferrin receptor (TfR). Input in A is 1/20^th^ of the total sample. (B) Input from Figure 8 blots (same experiment) to show LYN and pSFK input amounts. n.s. = nonspecific band.

There was dramatically more FYN, and activated FYN recovered by pSFK IP, in PAG1™^-^ endosomes in serum-free conditions compared to +serum (Figure 9A, FYN, pSFK). In WT cells, with serum, there were lower amounts of sequestered total FYN and LYN and active SFKs (Figure 8A). The data suggest that with low growth factor signaling (-serum) active SFKs were sequestered in MVBs and thus separated from interacting proteins in the cytosol (Figure 8A). In cells lacking functional PAG1, this pattern was disrupted, and both active and inactive SFKs were sequestered in MVBs. SFKs were constitutively active in PAG1™^-^ cells in serum-starved conditions (Figure 1E), and more were detected in MVBs (Figure 8 and 9).

Together, the data suggest that sequestration into the lumen of MVBs is a newly-identified mechanism to control SFK activity. It was previously known that PAG1 regulates SFK activity by serving as a scaffold to bring SFKs together with proteins that regulate them (e.g., CSK) (Oneyama et al., 2008). Our data suggest that PAG1 also directs SFK localization to the plasma membrane or endosomes depending on growth factor signaling. Furthermore, the selective ablation of signals that induce differentiation (NGF, GDNF; Figures 7, S1) but not proliferation (EGF; Figure 5) caused by expression of a truncated PAG1 suggest that this mechanism is responsible for distinguishing responses to different RTKs.

## Discussion

We had previously shown that the SFKs, FYN and LYN, changed intracellular location in endosomes and lipid rafts in response to stimulation of different RTKs (Palacios-Moreno et al., 2015). In addition, phosphoproteomics identified that the SFK scaffold protein, PAG1, enriched and highly phosphorylated in endosomes and lipid rafts, frequently colocalized with FYN and LYN in neuroblastoma cells (Palacios-Moreno et al., 2015). Specifically, signaling by the receptors ALK and KIT increased the amount of FYN and LYN in cellular fractions containing late endosomes and multivesicular bodies. In contrast, signaling by TRKA and RET, two receptors known to induce neuronal differentiation in sympathoadrenal neural crest lineages and neuroblastoma cell lines, shifted FYN and LYN from late endosome/MVB fractions to plasma membrane lipid rafts (Palacios-Moreno et al., 2015). We hypothesized that PAG1 may play a role in this shiftable localization of SFKs in response to RTKs to control downstream cell signaling effectors and responses. Here we demonstrated that expression of PAG1 lacking the transmembrane domain (PAG1™^-^) changed SFK activity and localization and abrogated cell differentiation signals but left proliferation signals intact.

Textbooks present the simplified view that RTKs all activate the same four canonical intracellular signaling pathways (RAS/ERK, PLC-γ, SFK, and PI3K/AKT), yet some RTKs cause proliferation, and others differentiation. Marshall (1995) proposed that sustained ERK signaling is required for differentiation. Chen et al. (2012) described a gradient of ERK vs. AKT activation that controls PC12 cell responses to NGF. In the latter model, higher levels of AKT activation lead to increased cell proliferation while higher levels of ERK activation lead to cell differentiation or lower rates of proliferation. Our data address cell signaling events that likely control ERK and AKT responses; we found that directed localization of SFKs to plasma membrane vs. the lumen of MVBs is an important mechanism to control differentiation. Sequestration in MVBs in PAG1™^-^ cells likely abrogates SFK activation at the plasma membrane in response to differentiation signals, and therefore decreases subsequent cellular responses such as ERK activation (Figures 7, S1), and extension of neurites (Figure 4). We hypothesize that PAG1™^-^ acts as a dominant negative by binding CSK and other cytoplasmic SFK-interacting proteins, which may explain why the effect on differentiation we observe is stronger than that seen in PAG1 knockout mice (Lindquist et al., 2011). Conversely, an increase of active FYN and LYN in endosomes enhanced responses to EGFR and KIT receptors (Figures 5, 6), and led to increased in growth rate and anchorage independent growth in PAG1™^-^ cells (Figures 2, 3). Thus, our results suggest that precise spatiotemporal regulation of SFK activity distinguishes RTKs that promote differentiation from those that promote proliferation, and that PAG1 plays a role in this process, which extends the model that defines RTK signals for differentiation (Chen et al., 2012b; Marshall, 1995).

PAG1™^-^ cells exhibited increased proliferation (Figure 2), and increased number of cells in the S and G2/M stages of the cell cycle under differentiation conditions (Figure 4). In agreement with previous work showing that AKT activation occurs predominantly after DNA replication in G2/M stages of the cell cycle (Liu et al., 2014), we found that cells with high levels of pAKT were correlated with the number of cells in the G2/M stages of the cell cycle for both PAG1™^-^ and wild type cells (Figure S2). This explains why we detected less pAKT in neuroblastoma cells under long-term differentiation conditions where cell division was reduced (unpublished results). The pAKT response to EGF is more rapid than the pERK response, peaking at about 3 min (Zheng et al., 2013), which likely explains why significant changes in AKT activation in response to ligands at 5 and 60 min as for pERK (Figures 5-7) were difficult to detect using flow cytometry methods used here. PI3K and AKT signaling are important for coordinating proliferative growth factor signaling and metabolism (Bilanges et al., 2019; Manning and Toker, 2017), yet it may be that only low levels of AKT activity are required for neuronal differentiation. Nevertheless, given the interconnected nature of cell signaling pathways (Grimes et al., 2018; Zheng et al., 2013), we can predict coordination with other signaling pathways downstream of RTK activation during differentiation, such as PLC-γ, PI3K, AKT, and mTOR complexes, which are also involved in endocytic trafficking and lysosomal degradation (Bilanges et al., 2019; Manning and Toker, 2017). In any case, our results support a role for SFKs, namely FYN and LYN, and control of their activity and intracellular localization by PAG1, as a mechanism to distinguish differentiation signals caused by the RTKs TRKA and RET.

A defect in differentiation, together with increased activity of total SFKs we observed in cells expressing PAG1™^-^ (Figure 1) is consistent with previous observations that PAG1 acts as a tumor suppressor in neuroblastoma cells (Agarwal et al., 2016; Oneyama et al., 2008). Oneyama et al. (2008) first demonstrated the interaction between PAG1 and SRC. They found that PAG1 mediates the inhibitory C-terminal phosphorylation of SFKs via binding CSK, and that a lipid-raft anchored SRC variant was largely inactive. Consistent with our soft agar assay data (Figure 3), cells lacking PAG1 formed almost 20 times as many colonies as those re-expressing functional PAG1 in colony transformation assays (Oneyama et al., 2008). Using siRNA knockdown of PAG1, Agarwal et al. also demonstrated increased SFK activity, increased anchorage independent growth, and increased ERK activity in neuroblastoma cell lines (Agarwal et al., 2016). A PAG1 mutant that lacked residues required for lipid raft localization failed to suppress SRC-mediated transformation (Oneyama et al., 2008), which emphasizes the importance of membrane localization of PAG1 for regulation of SFK activity. These data are consistent with our results with a PAG1 mutant lacking the transmembrane domain. Together, these studies suggest that defects in PAG1, whether through mutation or change in protein amount, have pro-proliferative effects and solidify its role as a tumor-suppressor.

PAG1 binds several different SFK family members, and can bind to more than one at a time, as well as to the kinase CSK that phosphorylates the SFK inhibitory site (Good et al., 2011; Ingley, 2008; Okada, 2012). PAG1 contains two proline-rich domains that interact with SFK SH3 domains, and is a substrate for phosphorylation on multiple tyrosine residues by SFKs. These phosphorylated sites then bind to SFK SH2 domains, maintaining SFKs in the active state when bound, even if phosphorylated on the inhibitory site (Solheim et al., 2008b). PAG1 has been also implicated in SFK dephosphorylation and ubiquitination (Ingley, 2008). Thus, an array of proteins bound to the PAG1 scaffold may either positively or negatively regulate SFK activity. While it had been assumed that PAG1 acts as a tumor suppressor by mediating the inhibitory interaction between SFKs and CSK, the anti-transformative effect of PAG1 was only partially dependent on CSK binding (Oneyama et al., 2008; Resh, 2008). This suggests that some of PAG1’s tumor suppressive function could involve regulation of SFK activity through endocytosis and sequestration as elucidated in these studies. In addition, PAG1 mediates interactions with other signaling pathways. PAG1 interacts positively with other RTK effectors, such as PLC-γ and PI3K (Ingley, 2008). It also interacts with the ubiquitin ligase SOCS1, and the phosphatase PEP (PTPN22), that negatively regulate SFK signaling (Ingley, 2008). SFK localization may also be influenced by other scaffolding proteins. SFKs interact with other signaling scaffold proteins at the plasma membrane and on endosomes, such as DOK adaptors, that may affect their trafficking and activation (Davidson et al., 2016).

Sequestration of active SFKs into MVBs represents a previously unappreciated method by which cells control SFK activity (Figure 8). FYN and LYN have been seen in structures that appear to be endosomes by fluorescence microscopy (Donepudi and Resh, 2008), and previous work showing SRC trafficking to, and regulating several components of, the endo/lysosomal pathway predicts that they should be found in MVBs (Reinecke and Caplan, 2014). Our studies suggest that MVB sequestration regulates SFK kinase activity towards cytosolic substrates similar to GSK3β kinase sequestration in the WNT signaling pathway (Taelman et al., 2010; Vinyoles et al., 2014). The data suggest that in the absence of signaling, such as during serum starvation, active SFKs are sequestered in MVBs, and RTKs, especially those that induce differentiation, keep SFKs active at the plasma membrane in lipid rafts. This mechanism was disrupted in PAG1™^-^ cells, which sequestered large amounts of total FYN and LYN (Figure 8). Interestingly, only FYN was immunoprecipitated by anti-phospho-SFK antibodies in endosome fractions, and activated FYN dramatically increased in MVBs under serum starvation conditions (Figure 9). In contrast, LYN was not immunoprecipitated by anti-phospho-SFK antibodies in endosomal fractions (Figure 9). This implicates sequestration of active FYN as the primary defect that abrogates the differentiation response. Sequestration of active FYN apparently had no inhibitory effect on growth rate or colony formation, which explains insensitivity to PP2 in PAG1™^-^ cells (Figures 2, 3), since PP2 inhibits FYN more effectively than SRC and LYN (Brandvold et al., 2012; Hanke et al., 1996).

We previously identified a potentially antagonistic relationship between FYN and LYN activation using computational analysis to identify patterns in phosphoproteomic data (Palacios-Moreno et al., 2015). In many neuroblastoma cell lines, activated FYN (pY420) was frequently associated with inhibited LYN (pY508); and in other cases inhibited FYN (pY531) was frequently associated with several phosphorylations on PAG1 (Palacios-Moreno et al., 2015). These data suggest that an antagonistic relationship between FYN and LYN activation, potentially mediated by an inhibitory interaction between PAG1 and FYN, may distinguish signals from different RTKs. While SRC’s normal function centers around cell migration, adhesion, and receptor endocytosis (Yeatman, 2004), FYN and LYN appear to have specialized roles in differentiation. FYN and LYN are delivered to the plasma membrane by different mechanisms than SRC (Van ’t Hof and Resh, 1997; Sato et al., 2009). In contrast to singly-acylated SRC, FYN and LYN are dually acylated by myristoylation and palmitoylation, which enhances their functional localization with lipid rafts (Bijlmakers, 2009; de Diesbach et al., 2008; Resh, 1994). FYN has two alternately spliced isoforms that differ in expression, binding partners and regulation (Brignatz et al., 2009). While LYN is hyperactive in glioblastoma (Stettner et al., 2005), FYN has been shown to play a role in differentiation in neurons induced by neurotrophin receptors (Patel et al., 2000; Pereira and Chao, 2007); zebrafish eye (St. Clair et al., 2018); migrating neurons in mammalian cortex (An et al., 2014; Zhang et al., 2016); neurons derived from stem cells (Zhang et al., 2016); myelin formation from oligodendrocytes (Colognato et al., 2004; Hossain et al., 2010; Peckham et al., 2016; Simons and Nave, 2016); mammary cells (Zucchi et al., 2004), and osteoclasts (Kaabeche et al., 2004; Kim et al., 2010). Other relevant roles have been described for FYN in the epithelial-mesenchymal transition and metastasis (Gujral et al., 2014), cellular regulation of redox state (Noble et al., 2015), and apoptosis (Du et al., 2012). PAG1 protein expression is strongest in the brains of newborn mice, where it negatively regulates FYN and SRC by recruiting CSK into lipid rafts, causing phosphorylation of their inhibitory tyrosines (Lindquist et al., 2011). In adult mouse brain, PAG1 is associated with FYN and SRC, but less so with CSK, and SFK kinase activity is regulated differently than in newborn mice (Lindquist et al., 2011). These studies together with our observations that FYN activity and localization was more disrupted than LYN in PAG1™^-^ cells, which had a defect in neuronal differentiation, lead to the hypothesis that sustained activation of FYN at the plasma membrane in lipid rafts plays a primary role in neuronal differentiation. This is probably oversimplified and further work will be required to test this hypothesis and untangle the relationship between FYN and LYN and their activity in lipid rafts and the endocytic pathway.

Previous work shows that FYN and LYN are degraded in proteasomes, mediated by CBL (Ghosh et al., 2004; Kaabeche et al., 2004). Our phosphoproteomic data indicated that CBLB was one of the most highly phosphorylated proteins in endosomes, while CBL was predominately phosphorylated in plasma membrane and lipid raft fractions (Palacios-Moreno et al., 2015). Both CBL and CBLB have proline-rich domains that bind to SH3 domains of several proteins, and ubiquitin associated (UBA) domains that mediate their function as E3 ubiquitin ligases (Fang et al., 2001; Kowanetz et al., 2003). These results, and other data showing that CBLB is uniquely associated with trafficking of the B cell antigen receptor (BCR) to late endosomes (Veselits et al., 2014), suggests a role for CBLB in ubiquitination of endosome-associated proteins, including FYN and LYN, making them available for inclusion as cargo into luminal vesicles in MVBs by the ESCRT pathway (Vietri et al., 2020).

Sequestration into MVBs may simply lead to protein degradation in the lysosome; however, there appear to be different populations of MVBs with longer half-lives, which may give rise to additional signaling functions (Huotari and Helenius, 2011). Receptor signaling from endosomal membranes after endocytosis influences temporal and spatial regulation of signaling effector pathways downstream from receptor activation (Bergeron et al., 2016; Irannejad et al., 2015). The presence of activated FYN in the lumen of MVBs, together with its role in neuronal differentiation noted above, suggests that FYN is potentially another important component of signaling endosomes in neurons, which play an important role in nervous system development and neurodegenerative disease (Bergeron et al., 2016; Chen and Mobley, 2019; Cosker and Segal, 2014; Harrington and Ginty, 2013; Marlin and Li, 2015; Scott-Solomon and Kuruvilla, 2018). Signaling endosomes containing NGF and TRKA have been identified in neurons to be Rab7 positive MVBs (Von Bartheld and Altick, 2011; Harrington and Ginty, 2013). Back fusion of luminal vesicles from MVBs would release active FYN and other kinases identified in signaling endosomes into neuronal cell bodies after retrograde transport (Von Bartheld and Altick, 2011; Bissig and Gruenberg, 2014; Falguières et al., 2012; Ye et al., 2018). Strong evidence for this was obtained by Ye et al., who showed that TRKA-containing MVBs are retrogradely transported in neurons and persist in the cell body, where TRKA is then re-expressed on the surface of endosomes and thus has access to cytosolic substrates (Ye et al., 2018). Further work will be necessary to determine whether FYN (and other SFKs) are retrogradely transported in the lumen of axonal MVBs and subsequently released into neuronal cell bodies.

MVBs can also fuse with the plasma membrane and release intraluminal vesicles as exosomes. Exosomes containing Wnt10b promote axon regeneration after neural injury (Tassew et al., 2017), and it is possible that “signaling exosomes” also play a role in nervous system development. We detected FYN and LYN in exosomes isolated from WT and PAG1™^-^ cell culture media (Figure 8B). Release of active FYN and LYN in exosomes, which can bind and signal to neighboring cells, may be one method by which global pSFK signals are increased in PAG1™^-^ cells (Figure 1). In accordance with this idea, more FYN was detected in PAG1™^-^ exosomes, and more FYN was immunoprecipitated with pSFK in endosomes (Figures 8, 9).

In conclusion, the data presented here further support a role for PAG1 as a key integrator of RTK signaling downstream of receptor activation and support its previously identified role as a tumor suppressor. Our data, together with our previous findings (Palacios-Moreno et al., 2015), fit with a model where PAG1 directs SFK intracellular localization to plasma membrane or endosomal compartments depending on the particular receptor activated by its ligand. PAG1 thus connects upstream RTK signals to SFKs, which in turn control other downstream effectors to induce cell differentiation. Furthermore, we demonstrate a new mechanism for regulation of SFK activity: sequestration into multivesicular bodies, which impacts many aspects of SFK signaling. These results support the broad hypothesis that the spatial regulation of pathways downstream of RTK activation within endosomes is a key mechanism by which cells determine a response to extracellular signals (Bergeron et al., 2016).

## Supporting information

Manuscript text with figures

## Acknowledgements

This work was supported by NIH NS070746-01, NS061303-01, COBRE NCRR Grant P20 RR015583 and generous donations from Dr. Craig Wilkinson. L.F. was supported by COBRE NCRR Grant P20GM103546.

## Author Contributions

Conceptualization, L.F. and M.G.; Methodology, L.F., J.P.M., M.G.; Investigation, all authors; Resources, T.L.; Writing – Original Draft L.F., M.G.; Writing – Review and Editing, L.F., M.G.; Visualization L.F., M.G.

## Materials and Methods

### Cell Culture

SH-SY5Y neuroblastoma cell lines were cultured in RPMI 1640 medium (Thermo Scientific HyClone, U.S.) supplemented with sodium bicarbonate (Sigma, U.S.) and 10% fetal bovine serum (FBS, Corning). Cells were maintained in a humidified incubator at 37°C, 5% CO_2_.

### CRISPR/Cas9 targeting of PAG1

CRISPR/Cas9 plasmids and sgRNAs targeting Exon 5 and 7 of the PAG1 gene were provided by Blake Wiedenheft and Royce Wilkinson (Montana State University). sgRNA sequences were housed in the Lentiviral Cas9/sgRNA vector (Addgene #57828).

Exon 5 sgRNA: 5’-GAAGCCGCGACAGCATAGTG GGG-3’.

Exon 7 sgRNA: 5’-GCAGATCCGAGGCCGATGTC TGG-3’.

The plasmid carrying Cas9/sgRNA was co-transfected into HEK293FT cells with the lentiviral packaging vectors psPAX2 (Addgene #12260) and VSV-g (Addgene #8454) following the Lipofectamine 3000 protocol (ThermoFisher) to generate lentivirus. Supernatants containing lentivirus were filtered through a low-protein binding syringe filter (0.45 μm, Millipore #SLHP033RS) before transductions. SH-SY5Y cells were transduced with different dilutions of virus-containing supernatant, and transfected cells were selected for by Puromycin (3 μg/mL) addition to culture media. Pools of puromycin-resistant cells were analyzed by western blot for PAG1 expression, and those lacking PAG1 were characterized further. DNA surrounding the target site was amplified by PCR, PCR products were sequenced by Eurofins. While PAG1 expression appeared to be ablated in initial screens, we subsequently noted a truncated, soluble protein expressed in these cells (described in Results).

### Neurite Extension

Cells were seeded according to calculated doubling times to control for initial increases in proliferation. Wild type SH-SY5Y (30,000 cells/well) or PAG1™^-^ SH-SY5Y (9,000 cells/well) cells were seeded in 6-well plates containing RPMI 1640 + 10% FBS. Cells were grown for 2-4 days until fully adhered. Spent media was aspirated and replaced with differentiation media composed of RPMI 1640 + 2% FBS, 10 µM 9-cis-Retinoic Acid (Sigma-Aldrich) and/or 5 nM human β-NGF (Peprotech). Cells were grown in differentiation media for 8 days; treatment media was replaced every other day. On day 8, neurite tracings, length quantifications, and cell body measurements were performed using ImageJ (Rueden et al., 2017). Only neurites longer than one cell body length were quantified to eliminate lamellipodia/filopodia. Representative images were taken on day 8 using a Zeiss Invertoskop light microscope with an Amscope MD500 camera.

### Mass Spectrometry

Tandem mas tag mass spectrometry was performed on SH-SY5Y and PAG1™^-^ cells as previously described (Grimes et al., 2018). Briefly, cells were washed and harvested in PBS and cell pellets frozen in liquid nitrogen. Cells were lysed in a 10:1 (vol/wt) volume of lysis buffer (4% SDS; 100 mM NaCl; 20 mM HEPES pH 8.5; 5 mM DTT; 2.5 mM sodium pyrophosphate; 1 mM β-glycerophosphate; 1 mM Na_3_VO_4_; 1 μg/ml leupeptin), and proteins were reduced at 60°C for 45 min. Proteins were then alkylated by the addition of 10 mM iodoacetamide (Sigma) for 15 min. at room temperature in the dark, and methanol/chloroform precipitated. Protein pellets were resuspended in urea lysis buffer (8M urea; 20 mM HEPES pH 8.0; 1 mM sodium orthovanadate; 2.5 mM sodium pyrophosphate; 1 mM β-glycerophosphate) and sonicated. Insoluble material was removed by centrifugation 10,000 xg, 5 min, and the supernatant diluted fourfold in 20 mM HEPES pH 8.5, 1 mM CaCl_2_, for Lys-C digestion overnight at 37°C, then diluted two-fold and trypsin digested 4-6 hours at 37°C. Samples were then acidified to pH 2-3 with formic acid. Peptides were purified on a Waters Sep-Pak column, dried in a speed-vac, and quantified using a micro-BCA assay (Thermo). Mass tag (10-plex TMT reagents; Thermo) were crosslinked to peptides in 30% acetonitrile/200 mM HEPES pH 8.5 1 hour at room temperature and the reaction stopped by the addition of 0.3% (v/v) hydroxyamine. Samples were then mixed in equimolar ratios, and the ratios checked and samples run on an Orbitrap Fusion Lumos MS (Thermo Fisher). Identification of peptides and quantification of mass tags was obtained from the from the MS2 spectrum after fragmentation by MS/MS analysis as described (Guo et al., 2014; Stokes et al., 2012). Peptides with False Discovery Rate (FDR) < 1% were selected for further analysis.

### Organelle Fractionations

Wild type LAN-6 or SH-SY5Y cells expressing either wild type PAG1 or PAG1™^-^ were grown to ∼90% confluency and serum starved for two hours prior to cell harvesting. Cells were then consecutively washed with cold PEE (1 mM EDTA, 1 mM EGTA, in 1X PBS) and PGB (0.1% glucose, 0.1% BSA, in 1X PBS) buffers, and resuspended in cold PGB containing the indicated RTK ligands. Cells were rotated with ligand for one hour at 4°C, washed with PGB, resuspended in cold PGB, and incubated for one hour at 37°C to permit endosomal trafficking. After internalization, cells were quenched in ice water, washed once each with PEE and Bud Buffer (38 mM aspartic acid, 38 mM glutamic acid, 38 mM gluconic acid, 20 mM MOPS pH 7.1 at 37°C, 10 mM potassium bicarbonate, 0.5 mM magnesium carbonate, 1 mM EDTA, 1 mM EGTA), and resuspended in Bud Buffer containing protease inhibitors. Cells were then mechanically permeabilized by a single passage through a Balch homogenizer. The “cracked” cells were centrifuged (1,000xg, 4°C, 10 minutes) to separate dense membranous fractions from organelles. The membrane pellet was further separated into detergent resistant membranes (DRM) and detergent soluble membranes (P1M) by addition of a mild detergent (0.1% IGEPAL) before centrifugation. The organelle-containing supernatant was layered on top of an iodixanol gradient (2.5-25% Optiprep, Sigma-Aldrich) and spun at 100,000xg to separate lysosomes (Lys), endosomes of high mass and density (E1), endosomes of intermediate mass and density (E2), endosomes of low mass and density (E3), and cytosol (Cyt) fractions. To isolate lipid rafts, an iodixanol gradient (2-30% Optiprep) was layered on top of DRM samples and spun to equilibrium at 100,000xg for 18 hrs. and floating and non-floating fractions were collected. Fractions from both types of gradients were taken using a Brandel fractionator. All centrifugations were performed using SW55Ti and MLS50 rotors in a Beckman ultracentrifuge. After fractions were collected from gradients, proteins were purified by TCA/acetone precipitation prior to suspension in 7 M Urea sample buffer for loading on SDS-PAGE.

### Protease Protection Assay

Fractions containing multivesicular bodies and late endosomes (labelled Endosomes) or early endosomes and cytosol (labelled Cytosol) were collected from organelle gradients after treatment (with or without serum starvation (2 hrs.)) according to the fractionation protocol. Bottom and top fractions were thoroughly mixed and divided into five equal volumes. Cold TCA (14%) was immediately added to the control fraction, the other four fractions were incubated at 37°C for either five minutes or thirty minutes with combinations of Proteinase K (0.01 µg/mL, Sigma-Aldrich) and/or IGEPAL (0.1%). Cold TCA (14%) was added after each incubation, and samples were stored at 4°C. Proteins in fractions were concentrated using TCA/acetone precipitation before SDS-PAGE and blotting on nitrocellulose.

### Immunoprecipitations

Following the organelle fractionation protocol with or without serum starvation (2 hrs.), cells were washed with cold PEE, suspended in Bud Buffer, and cracked using a Balch homogenizer. Fractions obtained after organelle fractionation were thoroughly mixed and divided into equal volumes before adding beads and/or primary antibody. Primary antibody and 10X CST Cell Lysis Buffer (Cell Signaling Technology, to 1X final concentration) were added to each IP sample. Samples were incubated, rotating, at 4°C overnight. A 10% bead solution (Protein A/G UltraLink Resin, Thermo Fisher #53132) was prepared in block buffer (1X lysis buffer containing 5% BSA). Beads were rotated 1 hr. at 4°C and washed twice with block. Beads were resuspended in 1 mg/mL BSA in 1X lysis buffer. Bead solution was added to each sample and incubated, rotating, at 4°C for 2 hrs. After incubation, IP beads were washed four times with 1X lysis buffer containing decreasing amounts of BSA (two washes with 0.25 mg/mL, followed by 0.1 mg/mL, and finally no BSA). Samples were resuspended in 7 M Urea sample buffer with DTT before SDS-PAGE and blotted on nitrocellulose.

### Flow Cytometry

Wild type SH-SY5Y and PAG1™^-^ SH-SY5Y cells were grown to 90% confluency before stimulation with RTK ligands. After seeding, cells were incubated for 24-48 hrs. prior to treatment to allow for attachment to the dish. Cells were harvested and washed with cold 1X PEE buffer before fixing in 4% paraformaldehyde in 1X PBS. After fixing, cells were treated with Benzonase in FACS buffer for 10 minutes at room temperature (to reduce cell clumping) prior to permeabilization with ice cold methanol. Samples were then incubated with Hoescht 33342 and/or the indicated fluorescent antibodies. Fluorescence minus one (FMO) controls were prepared from the control conditions within each experiment. Samples were read on a NxT Acoustic Focusing Cytometer (Life Technologies, Carlsbad, CA) using the Attune NxT Software, v2.4, and the resulting data were analyzed using Flowjo v10.0.

### MTT assay

SH-SY5Y WT cells were seeded at 50,000 cells/well and SH-SY5Y PAG1™^-^ cells were seeded at 5,000 cells/well in RPMI 1640 containing 10% FBS in a 96 well plate and allowed to adhere for 24 hrs. Spent media was aspirated and replaced with 100 µL of treatment media (0-5 μM PP2) and incubated for 48 hrs. before MTT addition. For time zero measurements, spent media was aspirated and media containing 5 mg/mL MTT was added and incubated for 4 hrs. After incubation, 40% w/v SDS in HCl pH 2 was added to dissolve MTT crystals. Absorbance was read at 570 nm in a VersaMax plate reader (Molecular Devices). Growth rates, defined as the number of doublings per day, were calculated by: R = [log_2_ (A_t_/A_0_)]/t, where A_0_ and A_t_ are absorbance measurements before and after treatment, respectively, and t is the number of days in culture. The growth rate index (GRI) was calculated as described by Hafner et al., by the formula: GRI=2^(R’/R)^-1 where R’ is the growth rate of cells under the test condition, and R is the control growth rate. In the case of PAG1™^-^ GRIs, R and R’ are the growth rates of WT and PAG1™^-^ cells respectively. For PP2 GRIs, R is the growth rate of cells in vehicle-control medium, and R’ is the growth rate of cells in PP2 medium. These calculations were carried out for five experiments. The resulting GRIs were pooled, and the second and third quartiles were used for analysis by plotting and in Welch’s t-tests. One-sample t-tests were also calculated for PP2 GRIs, to determine the statistical significance of growth rate inhibition (as indicated by deviation from 1). In addition, growth rates from all experiments were pooled, and Student’s t-tests were calculated to compare WT and PAG1™^-^ cells.

### Soft agar assays

2-hydroxyethyl agarose (0.6%) was added to the bottom of each well in a six-well plate. Plates were incubated at 4°C for one hour to solidify the agar. After one hour, plates were incubated at 37°C for 30 minutes before use. SH-SY5Y WT and SH-SY5Y PAG1™^-^ cells were diluted to a concentration of 80,000 cells/mL. Pre-warmed pipettes were used to mix 3% agarose with growth media containing 10% FBS to a final concentration of 0.3% agarose. SH-SY5Y WT or SH-SY5Y PAG1™^-^ cells were incubated in the media/agarose solution containing drug (5 µM PP2). The cell-containing solutions were then mixed with 2 mL of 0.6% agarose. The cell-agarose solution was added on top of the first agarose layer and incubated at 4°C for 15 minutes to solidify the agar. Cells were then incubated at 37°C for one week. After one week, the number of colonies over 70 µm in diameter were counted at 40X magnification. A grid was placed over each well which divided the well into six sections and one field of view was counted for each section. The average number of colonies per well and the average number of colonies per condition (average of three wells) was calculated.

### Exosome Isolation

Exosomes in Figure 8 were isolated using the ME Kit for Exosome Isolation (New England Peptide). WT SH-SY5Ys or PAG1™^-^ SH-SY5Ys were serum-starved for 16 hrs. before collecting cell culture media to prevent contamination from exosomes in fetal bovine serum. Ultracentrifugation was also used to verify the presence of LYN and FYN in exosomes. WT SH-SY5Ys or PAG1™^-^ SH-SY5Ys were serum-starved for 16 hrs. Cell culture media was harvested and centrifugations were performed to eliminate contamination from apoptotic bodies: 2,000xg, 20 min., 4°C; 5,000xg, 60 min. 4°C. The supernatant was collected and centrifuged at 100,000xg, 90min., 4°C; the exosome pellet was resuspended in 7M Urea sample buffer for SDS-PAGE and western blotting.

### Antibodies

#### Flow Cytometry

Anti-Human/Mouse phospho-SRC (Y418) PerCP-eFluor 710 (Affymetrix/eBioscience); AF647 Mouse anti-Src (pY418) (BD Biosciences); Hoechst 33342 (Cell Signaling Technology (CST) #4082); β-III Tubulin (CST #4466); pAKT-PE (pT308) (BD Biosciences); Rabbit anti-pERK-AF488 (Phospho-p44/42 MAPK (Erk1/2) (Thr202/Tyr204)) (CST #13214); Mouse IgG2b K Isotype Control PerCP-eFluor 710 (Affymetrix/eBioscience); Rabbit IgG Isotype Control AF488 (CST #4340); PE Mouse IgG1, κ Isotype Control (BioLegend #400111); AF647 Mouse IgG1, κ Isotype Control (BioLegend #400135).

#### Western Blots

Fyn Antibody 1:1000 (CST #4023); Lyn Antibody 1:1000 (CST #2796); PAG1 (Csk-binding protein antibody PAG-C1) 1:1000 (ThermoFisher #MA1-19287); Phospho-Src family (Tyr416) 1:1000 (CST #6943/2101); Src 1:1000 (CST #2109); Transferrin Receptor (TfR) H68.4 1:7500 (ThermoFisher 13-6800); Flotillin 1:1000 (CST #3253); ECL Anti-Mouse IgG HRP linked secondary 1:5000 (GE Healthcare #NA931); ECL Anti-Rabbit IgG HRP linked secondary 1:5000 (GE Healthcare #NA934).

### Ligands and Inhibitors

PP2 (Calbiochem #529573); 9-cis-Retinoic Acid (Sigma Aldrich # R4643); EGF – 5nM (PeproTech #AF-100-15) (R&D #236-EG-200); GDNF (PeproTech #450-10) (R&D #212-GD-010); β-NGF (PeproTech #450-01) (R&D #256-GF-100); SCF (PeproTech #300-07) (R&D #255-SC-200); PTN (R&D #252-PL).

