## Supplementary material for "PAG1 directs SRC-family kinase intracellular localization to mediate receptor tyrosine kinase-induced differentiation": Manuscript text with figures

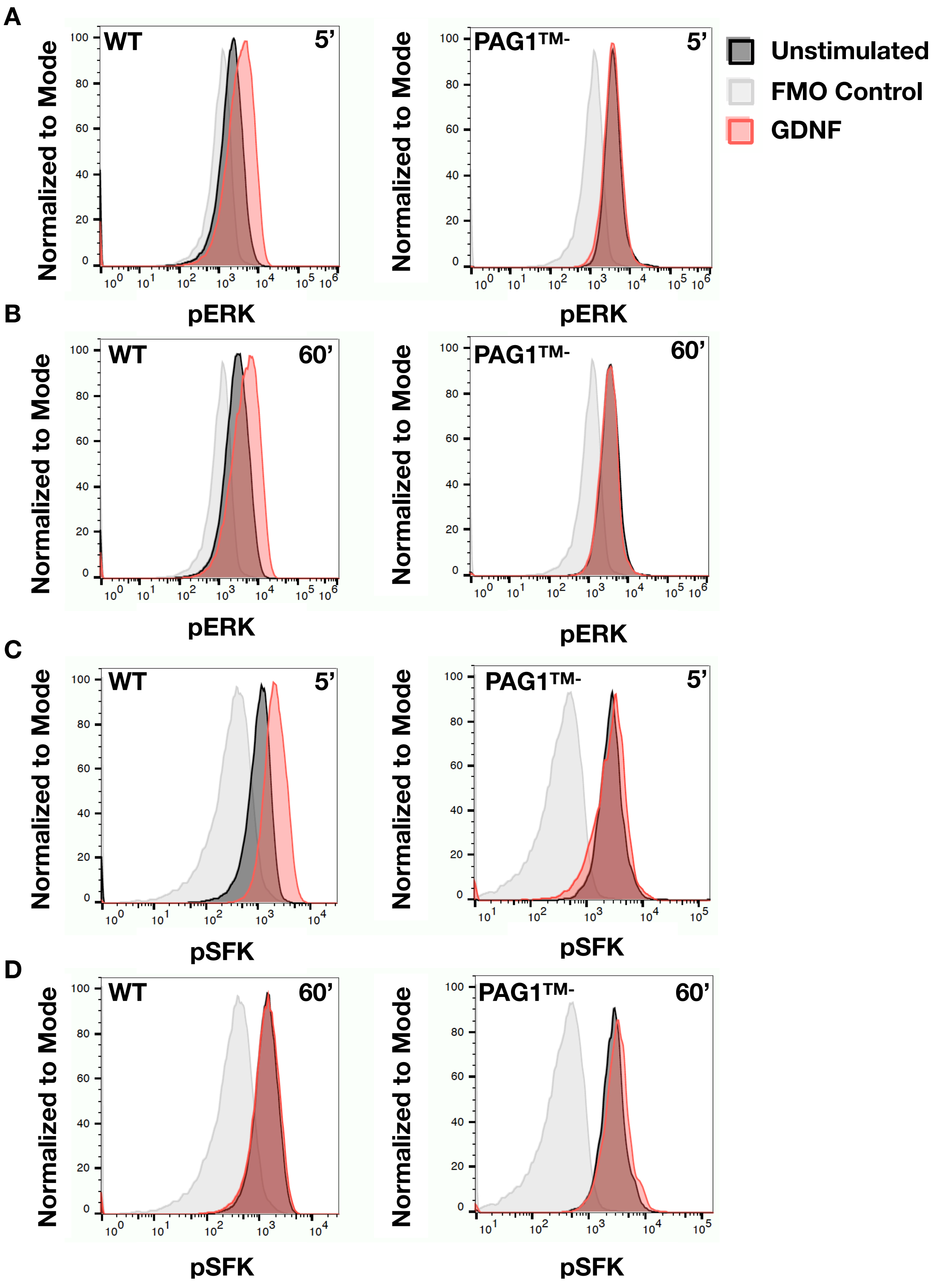


**Supplementary Figure 1: PAG1TM- decreased responses to pro-differentiation receptor RET**. Flow cytometry analysis of pERK (A, B) and pSFK (C, D) activation after exposure to 5nM GDNF for either 5 minutes (A and C) or 60 minutes (B and D). Adherent SH-SY5Y cells expressing either wild-type (WT) PAG1 or PAG1TM- were treated with ligand before fixing and staining with fluorescent antibodies for pERK (Phospho-p44/42 MAPK (Erk1/2) (Thr202/Tyr204)) and pSFKs (pY416). Data displayed are from one experiment representative of at least three independent experiments. Cell counts were normalized to mode to account for differences in final cell number.

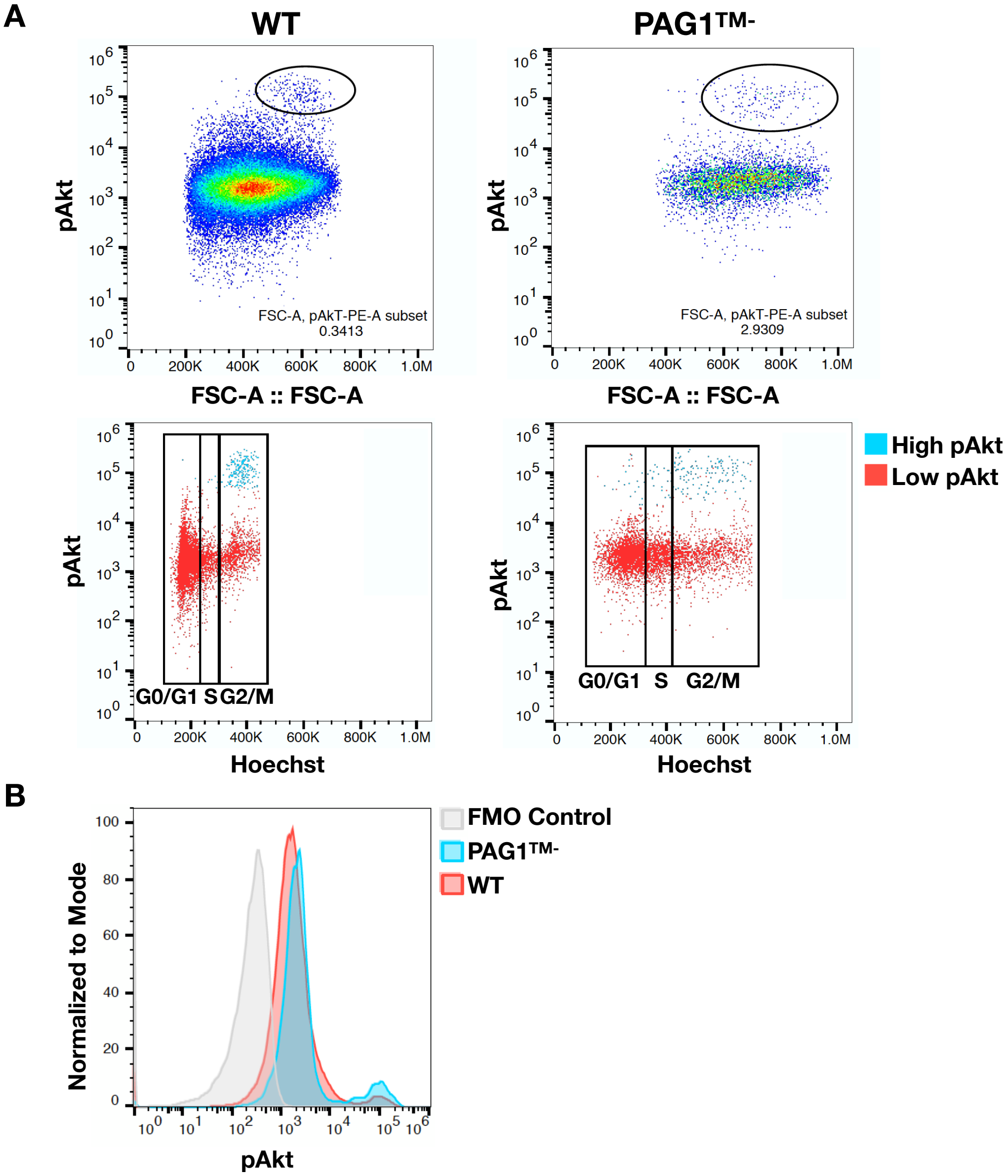


**Supplementary Figure 2: Akt activation in WT and PAG1TM- cells correlates with cell cycle stage G2/M.** Flow cytometry analysis of pAkt (pT308) and DNA content (Hoescht). (A) Top panels compare cell size (FSC) to pAkt signal. Cells with high pAkt (circled) are highlighted in blue in bottom panels comparing DNA content to pAkt signal. Cell cycle stages determined by DNA content are demarcated by boxes. (B) Comparison of pAkt signal between WT and PAG1TM- cells, cell counts were normalized to mode on the y-axis to account for differences in final cell number.

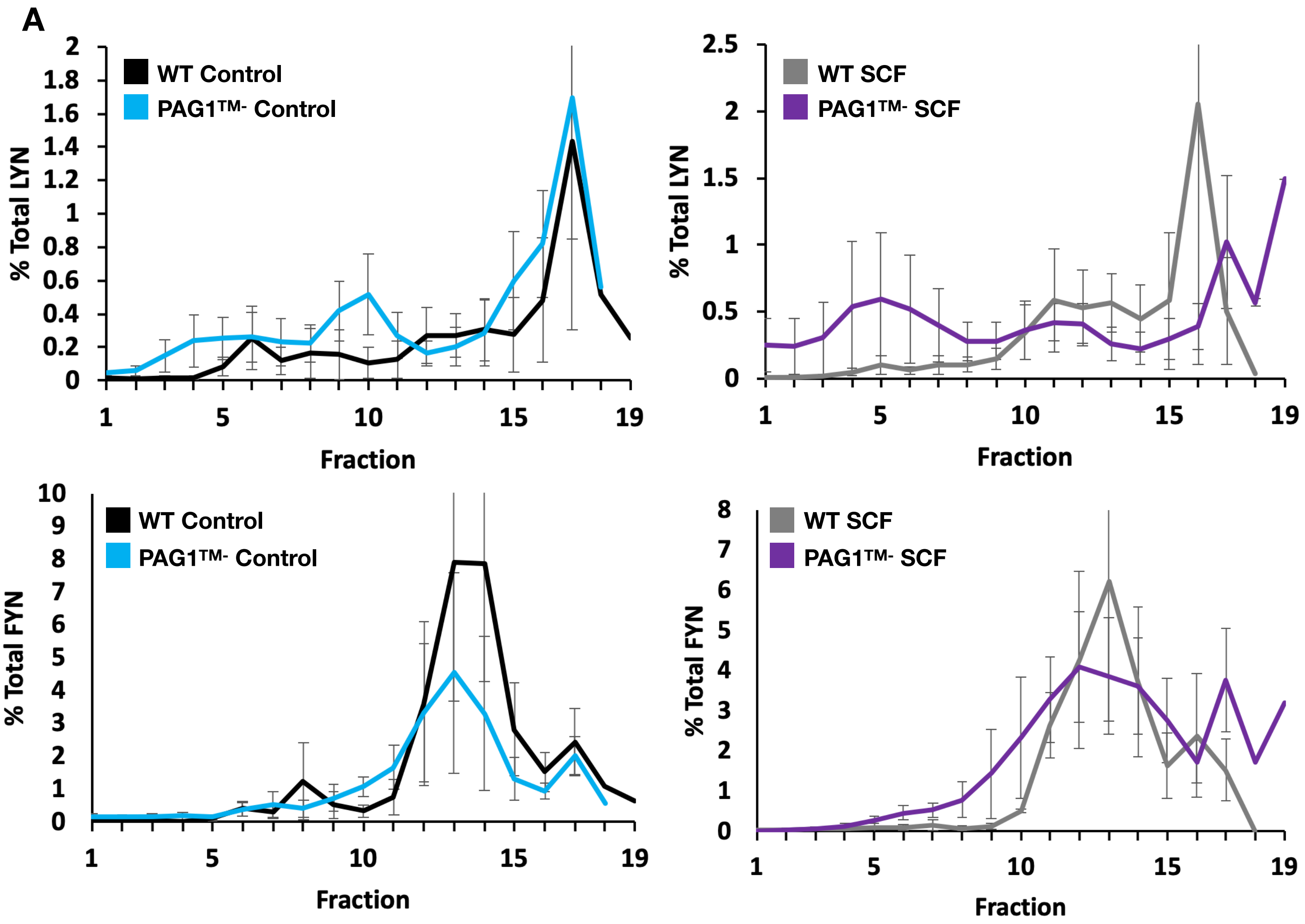


**Supplementary Figure 3: FYN and LYN distributed into different organelles.** Organelles were harvested from WT and PAG1TM- SH-SY5Ys after Control or SCF (5nM) treatments. Cell fractions containing organelles were separated by gradient ultracentrifugation. Fractions decrease in density, with fraction 1 being the densest and fraction 20 being the least dense. Following centrifugation, fractions were run on western blots and the amount of total LYN (A) or FYN (B) was quantified as the % of signal in each fraction vs. the total amount of signal.
